# Plant-Compatible Xenium *In Situ* Sequencing: Optimised Protocol for Spatial Transcriptomics in *Medicago truncatula* Roots and Nodules

**DOI:** 10.1101/2025.07.22.663073

**Authors:** Min-Yao Jhu, Jo Heffer, Alex Deamer, Thiago Alexandre Moraes, Ania M. Piskorz, Chongjing Xia

## Abstract

Elucidating the spatial and temporal regulation of gene expression during plant organogenesis is crucial for enabling precise crop improvement strategies that incorporate beneficial traits into crops while avoiding adverse effects. Root nodules, specialised organs formed in symbiosis with nitrogen-fixing bacteria, provide a valuable system to study cell-type-specific gene networks in a symbiosis-induced developmental context. However, capturing these dynamics at cellular resolution in intact plant tissues remains technically challenging. Spatial transcriptomics technologies developed for animal systems are often not directly transferable to plant tissues due to fundamental differences in tissue composition between plants and animals, including rigid and heterogeneous plant cell walls, high cell wall autofluorescence, and large vacuoles in plant cells that complicate probe access and signal detection.

To address these challenges, we present an optimised protocol for applying the Xenium *in situ* sequencing platform to formalin-fixed paraffin-embedded (FFPE) sections of plant tissues, including *Medicago truncatula* roots and nodules. Key technical adaptations include customised tissue preparation, optimised section thickness, hybridisation conditions, post-Xenium staining, imaging, and downstream image analysis, all tailored specifically for plant samples. To mitigate autofluorescence and enhance detection sensitivity, we employed a strategic approach to codeword selection during probe design. Furthermore, we developed a modular probe design approach combining a custom 380-gene standalone panel with a 100-gene add-on panel. This design allows flexibility for addressing diverse research questions and includes orthologous gene sequences from two *Medicago* ecotypes, ensuring compatibility for downstream functional validation using mutant lines available in both genetic backgrounds. We validated the protocol across nodules at multiple developmental stages using both the 50-gene panel targeting mature nodule cell identity and the extended 480-gene panel, which includes markers across different cell types and developmental stages, as well as genes of interest identified from prior single-cell and bulk RNA-seq analyses. This optimised workflow provides a reproducible and scalable method for high-resolution spatial transcriptomics in plant tissues, establishing a robust foundation for adaptation to other plant species and developmental systems.

## 1 Introduction

Certain plants have developed extraordinary adaptive traits that allow them to thrive under conditions such as nutrient deficiency, notably through the ability to fix atmospheric nitrogen (N2) via symbiosis with bacteria in root nodules. These capabilities present a substantial opportunity to boost agricultural productivity (*1*). However, unravelling the genetic foundations conferring these adaptive advantages has been challenging through traditional genetics or bulk RNA sequencing methods. This difficulty arises because the emergence of such traits often involves altering the regulatory mechanisms of genes that are otherwise conserved, leading to novel spatial or temporal gene expression patterns—a phenomenon observed across both the plant and animal kingdoms. For example, genes that typically play a role in lateral root development have been repurposed in some plants to facilitate the formation of nitrogen-fixing nodules (*2*).

The potential agricultural benefits of these adaptive traits are immense, but identifying the precise genetic mechanisms that underlie their development poses a formidable challenge. These traits often result from evolutionary changes that modify the developmental fate of particular cell types. For instance, cells in the cortex, endodermis, and pericycle can be reprogrammed from their original role in lateral root formation to create nitrogen-fixing nodules. Pinpointing the transcriptional differences in cell lineage that lead to such divergent developmental outcomes is crucial in uncovering how distinct organ structures and functionalities emerge. Traditional approaches to studying cell lineage regulation in plant development have primarily focused on analysing individual genes, using methods such as ethyl methanesulfonate (EMS) mutagenesis, *Medicago Tnt1* retrotransposon insertional mutagenesis, or, more recently, CRISPR knockouts. However, these methods, including bulk RNA sequencing, lack the resolution to observe changes in gene expression patterns at the cellular level. Creating a detailed atlas covering the organogenesis of *Medicago* nodules, which form a mutualistic symbiosis with nitrogen-fixing rhizobia, is essential for understanding the complex interplay of cell types within. Grasping the development and differentiation of various cell lineages into distinct cell types and functions is a pivotal initial step in deciphering the gene regulatory networks that drive the formation of this specialised organ.

Single-cell RNA-sequencing (scRNA-seq) has revolutionised plant research by allowing scientists to group cells into populations with similar molecular characteristics, facilitating a deeper understanding of cell types and their states (*3*). In contrast to bulk RNA-seq, scRNA-seq enhances sensitivity by capturing signals from rare or transient cell populations that would otherwise be diluted in population averages. As technology advances, offering higher throughput and resolution, it enables increasingly precise cell classification. However, because traditional scRNA-seq relies on dissociated tissues, it disrupts the spatial context essential for studying organogenesis, particularly when we encounter cell populations that fall outside our existing knowledge of plant cell histology and physiology. As discussed in previous studies (*4*), examining the molecular identity of these cell populations and their spatial arrangement within plant tissues is essential to fully comprehend their roles and interactions. One of the go-to methods for mapping the spatial distribution of cell population markers discovered through single-cell transcriptomics is using transgenic reporter lines, which express fluorescent proteins driven by the predicted promoters of target genes. Although effective for demonstrating individual gene promoter activity, this method struggles with complex tissues because cell identity often depends on the expression of multiple genes. Additionally, generating transgenic plants is time-consuming, and the artificial expression may not reflect true gene activity due to missing genomic context, like enhancer-promoter interactions. Similarly, while traditional *in situ* hybridisation offers a partial solution by addressing some limitations, it lacks the ability to analyse many genes simultaneously due to its low multiplexing capabilities. Hence, comprehensive spatial gene expression analysis, crucial for understanding cell functions and interactions, requires examining a broad spectrum of genes at the single-cell resolution.

The advent of spatial transcriptomics is set to overcome these obstacles by providing detailed insights into both the molecular composition and the physical locations of cells within complex tissues (*5, 6*). Spatial transcriptomics techniques, such as spatially barcoded arrays (*7–9*) and multiplexed fluorescence *in situ* hybridisation (FISH) (*4, 10*), enable the study of extensive gene sets along with their spatial contexts. This advancement is particularly significant in plant research, opening new possibilities for exploring the complex interplay of plant cells and tissues (*4, 7–10*). However, the task of investigating complex gene regulatory networks during plant organogenesis, within their native spatial contexts, remains challenging. Adapting spatial transcriptomics methods from animal to plant tissues is especially difficult due to the unique structure of plant cell walls and the presence of large vacuoles within plant cells, which make fresh-frozen cryosectioning, required by platforms like Visium, particularly challenging in plant tissues and often insufficient for achieving true cellular or subcellular resolution.

To overcome these difficulties, we present an optimised protocol for spatial transcriptomics in plants using the Xenium *in situ* platform (10x Genomics). Xenium is a highly sensitive, multiplexed, fluorescence-based *in situ* sequencing method capable of detecting hundreds to thousands of transcripts at subcellular resolution in formalin-fixed paraffin-embedded (FFPE) sections. While Xenium has been successfully applied in animal systems, this is the first detailed protocol tailored for use in plant tissues. Using *Medicago truncatula* root nodules—a complex, symbiosis-induced organ formed at the plant–bacteria interface—as a model, we introduce a suite of plant-specific optimisations, including modifications in probe design, tissue preparation, sectioning, hybridisation, imaging, and cell segmentation.

Plant cell walls exhibit strong autofluorescence, which varies in intensity and emission spectrum depending on species, tissue type, and cell wall composition. By analysing autofluorescence spectra and excluding channels most affected by it during codeword selection, we improved transcript detection accuracy. These optimisations increased the high-quality transcript detection rate from approximately 75% to 95%. Additionally, the heterogeneous texture of plant tissues, due to variable cell wall composition, the presence of large vacuoles, diverse cell shapes and sizes, and rigid structures such as xylem, complicates sectioning. We therefore optimised tissue processing and section thickness to minimise tissue detachment, tearing, and compression artefacts.

Moreover, traditional cell segmentation methods used in animal tissues often rely on DAPI nuclear staining with radial expansion, which assumes round or elliptical cell geometry. This approach is not well-suited to plant cells, which often have irregular or polygonal shapes. We found that leveraging the inherent autofluorescence of plant cell walls, captured through fluorescence lifetime imaging microscopy (FLIM) and tau gating on a confocal microscope, provides a robust boundary signal for segmentation. This method significantly improved segmentation quality in plant tissues, enabling more accurate spatial mapping of transcript data to individual cells.

We validated this protocol across multiple stages of nodule development using both a 50-gene panel enriched for mature nodule cell-type markers and a 480-gene panel composed of a 380-gene standalone panel with a 100-gene add-on module. These panels target a broad range of developmental regulators, cell identity markers, and candidate genes identified in prior bulk and single-cell transcriptomic studies. While demonstrated in nodules, the protocol is broadly applicable to other plant tissues and systems requiring high-resolution spatial gene expression analysis.

By enabling high-resolution, multiplexed spatial transcriptomics in plant FFPE sections, our optimised Xenium protocol fills a critical methodological gap. It provides a scalable and reproducible workflow for studying plant development and gene regulation in situ, opening new opportunities for fundamental research and translational applications in crop improvement.

## 2 Materials

### 2.1 Probe Design

#### 2.1.1 Software and Resources

- **10x Genomics Xenium Designer** Web-based tool used for probe design and panel submission.
- Reference Genome Information Transcriptome reference files for the target species, e.g., *Medicago truncatula* mRNA FASTA file.
- Gene Expression Datasets Single-cell RNA-seq data (ideally from the same species, tissue type, and developmental stage). Files should be in .h5 format for compatibility with expression filtering and probe design.
- Target Gene List A curated list of target genes, including both transcript IDs (must match the reference genome annotation) and gene names.

- *Ortholog Gene List:* If designing probes for use across species or ecotypes, identify orthologous genes in each genome and organise them into a unified gene list for panel design.

### 2.2 Tissue Fixation and Paraffin Block Preparation

#### 2.2.1 Reagents and Solutions

- 1× **PBS Buffer:** 10 mM phosphate, 137 mM NaCl, 2.7 mM KCl (pH 7.2–7.4)
- Fixative (4% paraformaldehyde)
- Glutaraldehyde (GA, 25%)
- Ethanol (10%, 30%, 50%, 70%, 90%, 100%)
- Histo-Clear II
- Paraplast Xtra (melted, 58–60 °C)
- MQ water (autoclaved and de-gassed)

#### 2.2.2 Equipment and Tools

- Glass vials
- Vacuum desiccator
- Embedding molds
- Heating plate (60 °C)
- Dissecting microscope
- Ice or cooling plate

### 2.3 Paraffin Sectioning and Section Slide Preparation

#### 2.3.1 Reagents and Solutions

- RNaseZap

#### 2.3.2 Equipment and Tools

- Microtome with disposable blades
- Sectioning implements - forceps, brush, slide rack, ice tray
- Sectioning trimming implements – razor blade, cutting mat, lamp
- Waterbath (45°C) filled with Milli-Q water
- Slides: Xenium slide (room temperature, remove from freezer 30 minutes before) and standard slides
- Microscope
- Oven (42°C)
- Desiccator

### 2.4 Probe Hybridization and Processing Xenium Slides

#### 2.4.1 Reagents and Solutions

- Deparaffinization reagents

- Xylene
- Ethanol (100%, 96%, 70%)
- Nuclease-free water
- RNaseAWAY
- Lint-free Laboratory Wipes
- Nuclease-free Water
- 10x PBS, pH 7.4
- Tween-20, 10%
- Urea, 8M
- TE Buffer, pH 8.0
- Ethanol, 100%
- DMSO, 100%
- KCL, 2M
- Isopropanol, 100%
- Xenium Thermocycler Adaptor V1
- Xenium Slides & Sample Prep Reagents V1 (2 slides, 2 rxns) PN-1000460

- Xenium Slide
- Xenium Cassette V1
- Xenium Cassette Lids
- Xenium FFPE Tissue Enhancer
- Perm Enzyme B
- Probe Hybridisation Buffer
- Post-Hybridisation Wash Buffer
- Ligation Buffer
- Ligation Enzyme A
- Ligation Enzyme B
- Amplification Mix
- Amplification Enzyme
- Autofluorescence Mix
- Reducing Agent B
- Nuclei Staining Buffer
- Xenium Decoding Consumables (1 run, 2 slides) PN-1000487

- Extraction Tip
- Pipette Tips
- Xenium Buffer Caps
- Xenium Objective Wetting Consumable
- #1 Deionised Water Bottle
- #2 Xenium Sample Wash Buffer A Bottle
- #3 Xenium Sample Wash Buffer B Bottle
- #4 Probe Removal Buffer Bottle
- Xenium Decoding Reagents (1 run, 2 slides) PN-1000461

- Xenium Decoding Reagent Module A
- Xenium Decoding Reagent Module B
- Xenium 380 gene Standalone Custom Probes
- Xenium 100 gene Add-on Custom Probes

#### 2.4.2 Equipment and Tools

- Oven (60°C)
- Reagent pots
- Xenium analyser
- <u>Xenium Compatible</u> Thermocycler (Bio-Rad C1000 Touch™ Thermal Cycler with 96-Deep Well Reaction Module)
- P1000 Pipette + Low Retention Tips
- P200 Pipette + Low Retention Tips
- P20 Pipette + Low Retention Tips
- Pipette Boy
- 50ml/ 10ml/ 5ml Stripettes
- 500ml Glass Bottle
- 500ml Measuring Cylinder
- 50ml/ 15ml Falcon Tubes
- 5ml/ 1.5ml Low-bind Eppendorf Tubes
- Xenium Adhesive Seals
- Scissors
- Forceps

### 2.5 Post-Xenium H&E Staining

#### 2.5.1 Reagents and Solutions

- Quencher removal solution
- Milli-Q water
- Harris haematoxylin
- Tap water
- 2% Acid alcohol (add 2M Hydrochloric Acid to 70% ethanol to create a 2% solution)
- Eosin
- Ethanol 50% + 0.1% Tween 20
- Ethanol 70% + 0.1% Tween 20
- Ethanol 100%
- Xylene
- Distrene Plasticiser Xylene (DPX)

#### 2.5.2 Equipment and Tools

- Weighing scale
- Reagent pots
- Slide rack
- Automated stainer (Leica ST5020)
- Automated coverslipper (Leica CV5030)

Note: Any automated stainer and coverslipper can be used; if none are available, staining and coverslipping can be done manually.

### 2.6 Post-Xenium Confocal Imaging

- Leica Stellaris 8 FALCON FLIM Microscope

### 2.7 Post-Xenium Image Segmentation and Transcript Realignment

**1. Software**:

- **Python environment and Cellpose:** Version 2.0 or later is recommended.
- **Python packages:** A Python environment (e.g., Python 3.8+) with standard scientific libraries such as:
- **tifffile**, **imagecodecs**: For reading and writing image files, including TIFF, OME-TIFF metadata, and other formats.
- **numpy**, **scipy**, **pyarrow**: For numerical operations.
- **scikit-image**, **opencv-python**: For image processing tasks (though Cellpose handles much internally).
- **matplotlib**: For data visualization tasks.
- **pandas**: For handling tabular data if manipulating image metadata or results.

**2. Hardware**

- **CPU**: Cellpose can run on standard CPUs but performs faster on GPUs.
- **GPU (Recommended)**: Use an NVIDIA GPU with CUDA support for efficient processing of large images or batches.
- **RAM**: 8–32 GB+ recommended, depending on image size.

## 3 Methods

### 3.1 Advanced Custom Probe Design for Xenium Panel

10x Genomics currently offers several pre-designed tissue or pathway specific Xenium panels compatible with human and mouse species. These include options for Xenium v1 chemistry (247–380 gene targets) and Xenium 5k chemistry (5,000 targets). For other species, such as plants, the current available solution is to design a custom standalone panel, with support for up to 480 genes.

For our target species, *Medicago truncatula*, we began by designing a standalone custom panel containing 50 target genes for our initial two Xenium slide experiments. This panel was tailored to include key marker genes for specific cell types. Based on insights from the initial results, we developed an expanded standalone custom panel consisting of 380 genes, along with a compatible 100-gene custom add-on panel, for a total of 480 genes. This tiered design enables us to retain a core set of cell–type–specific marker genes in the 380-gene panel, while also allowing flexibility for future experiments by adding new gene sets relevant to specific research questions through the 100-gene add-on panel.

#### 3.1.1 Target Gene Selection

1. **Define Experimental Objectives**: Select genes of interest based on your biological question (e.g., genes involved in nodule development). Prepare a gene list with gene IDs that correspond to the reference genome version used for probe design.
2. **Use Transcriptomic Resources**: Utilize single-cell RNA-seq data to identify cell-type marker genes. These markers can serve as internal quality controls to validate spatial transcriptomic results and allow cross-referencing with existing datasets.
3. **Exclude Extremely Highly Expressed Genes (EHEGs)**: As Xenium In Situ uses fluorescence-based imaging to detect transcripts, avoid over-representation of transcripts from EHEGs, which can cause optical crowding. Include EHEGs only if they are essential to your biological hypothesis.
4. **Expression Range Guidelines**: Ideally, target genes should:

- **Optimal Expression Range:** Target genes ideally have an average expression between 0.1–100 transcripts per cell.
- **High Expression Caution:** Genes with expression levels above 100 transcripts per cell may lead to optical crowding. However, if the full panel capacity is not being utilised and the gene has been successfully tested in a pilot run, a few highly expressed genes can be retained if they are critical for addressing specific biological questions.
- **Low Expression Considerations:** Based on our experience, genes with low expression (below 0.1 transcripts per cell) can still be detected by Xenium if they exhibit cell– type–specific expression patterns, as the platform has high sensitivity. However, if a gene is both lowly expressed and lacks cell type specificity, the detected signal may be difficult to interpret and less biologically meaningful.
5. **Prioritise Informative Targets:** Give preference to genes with cell-type- or state-specific expression, as these offer more spatial resolution and biological interpretability.
6. **Identifying Orthologs for Conserved Probe Design:** If designing probes for use across different species, varieties, or ecotypes, it is advisable to select conserved regions based on ortholog information to ensure optimal hybridisation performance. For *Medicago truncatula*, the A17 and R108 ecotypes are commonly used in research and mutant line development. To maintain probe compatibility across both ecotypes, we first identified target gene orthologs using OrthoFinder (version 2.5.5) (*11*) with default parameters. Protein sequences for R108 and A17 were downloaded from https://data.legumeinfo.org and https://medicago.toulouse.inra.fr, respectively. For genes with more than one predicted ortholog, BLASTn was used to compare mRNA sequences, and the ortholog with the highest nucleotide sequence similarity was selected for probe design. Where orthologs could not be confidently resolved, ecotype-specific probes were designed instead.

#### 3.1.2 Codeword optimisation

1. **Assess Autofluorescence:** Use a confocal or fluorescence microscope to image practice FFPE sections (before and after H&E staining). Check for strong cell wall autofluorescence specific to your species and tissue type.
2. **Adjust Fluorophore Channel Usage:** If strong autofluorescence is present, identify its dominant wavelength range and avoid assigning frequently used codeword channels in that range. Request custom channel allocation in your design notes if needed.
3. **Consider Pilot Panel:** If feasible, begin with a small pilot panel (e.g., 50 genes) to evaluate optical crowding, autofluorescence impact, and detection sensitivity. Use these insights to refine your design before scaling up to a full panel.
4. **Plan for Optical Crowding:** As panel size increases, consider the combined effects of autofluorescence, tissue structure, and transcript abundance. Strategically distribute EHEGs and avoid clustering highly expressed targets within the same channel group.

#### 3.1.3 Panel Submission and Review

1. **Format Gene List**: Use the 10x Genomics template for gene list submission. Ensure that gene IDs match the selected reference genome assembly.
2. **Submit Panel Design**: Upload your formatted gene list to the Xenium Design Portal. Indicate any special requests regarding codeword/channel usage based on autofluorescence analysis.
3. Review QC Report:

- Check the number of probe pairs designed per gene (ideally ≥8). Genes with fewer probes may have reduced detection efficiency.
- Examine the QC flags for predicted optical crowding, especially for EHEGs. Consider removing flagged genes unless they are essential to your hypothesis.
4. **Finalise and Order Panel**: Confirm your final gene list based on QC feedback. Panel manufacturing typically takes ∼4 weeks after submission.

### 3.2 Tissue Fixation and Paraffin Block Preparation

#### 3.2.1 Buffer Preparation

**Day 1: Preparing Fixative Solution**

1. To prepare 30 mL of fixative solution, add 80 μL of 1 N NaOH to 24 mL of water in a glass beaker. This step helps adjust the pH for dissolving paraformaldehyde (PFA).
2. Warm the solution slightly in a microwave. Heat for 10 seconds, repeating 4 times, and mix well after each round.
3. Add 1.2 g PFA to the warm solution. Stir continuously until the PFA fully dissolves. The solution may appear cloudy initially, but it will clear upon complete dissolution.
4. Cool the PFA solution on ice for approximately 15 minutes until it reaches 4°C before proceeding. Glutaraldehyde (GA) is heat-sensitive, and adding it to a warm solution may reduce the fixative’s effectiveness.
5. Add 300 μL GA (25%) and stir gently.
6. Finally, add 6 mL of 5× PBS to the mixture to achieve the desired buffer strength.
7. Dispense the fixative into clean glass vials and keep them on ice to maintain freshness.

#### 3.2.2 Harvesting and Tissue Fixation

**Day 1: Fixing Plant Samples**

1. Harvest *Medicago truncatula* nodules at the desired developmental stage. (Fig. 1)
2. Place plant tissues into the prepared fixative. Ensure the fixative volume is at least 10× the tissue volume for effective penetration.
3. Apply a vacuum (∼500 mm Hg or ∼0.065 MPa) to the samples while keeping them on ice. Hold the vacuum for 20 minutes, release it slowly, and repeat this process 4 times. This step removes air pockets, allowing the fixative to penetrate the tissues fully.
4. Incubate the samples at 4 °C overnight (at least 12–16 hours) to allow complete fixation. Notes: The vacuum time and strength might need to be adjusted depending on the samples that you are working with. Please observe the condition of your sample; a fully penetrated sample should sink to the bottom after the vacuum.

**Fig. 1.**
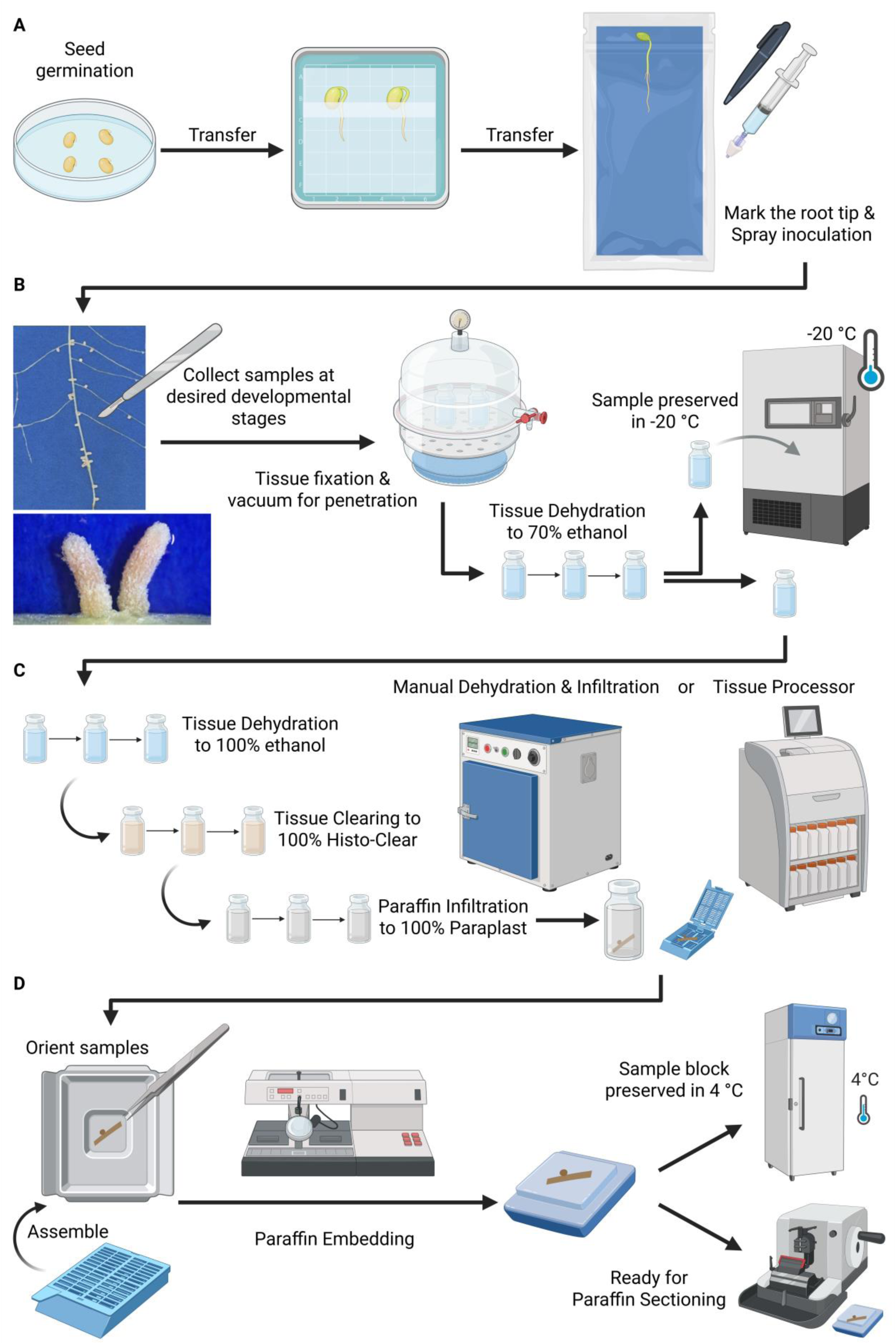
Workflow of tissue sample harvesting and sample block preparation. (A) Overview of tissue preparation: Start with *Medicago* seed germination and growth, followed by spray inoculation with rhizobia. (B) Harvesting and fixation: Excise desired tissue samples, fix in paraformaldehyde (PFA) solution under vacuum for effective penetration, followed by dehydration. Samples can be stored long-term in 70% ethanol at −20 °C until further processing. (C) Dehydration and paraffin infiltration: This chapter describes a primarily manual method for dehydration and paraffin infiltration, which is gentler and more easily optimised according to sample condition. While automated tissue processors are an alternative, a species- and tissue-specific optimised program is necessary. (D) Embedding: Fully paraffin-infiltrated samples are embedded by proper orientation in moulds, followed by histology cassette assembly and paraffin solidification. Solidified blocks can be stored at 4 °C until ready for paraffin sectioning.

#### 3.2.3 Tissue Dehydration

**Day 2: Dehydration**

1. Remove the fixative from the tissues using a pipette.
2. Pass the tissues through a graded ethanol series at room temperature (RT) to dehydrate them (Fig. 1):

- 10% ethanol: 30 minutes
- 30% ethanol: 30 minutes
- 50% ethanol: 30 minutes
- 70% ethanol: 30 minutes
- 90% ethanol: 30 minutes
- 100% ethanol: 1 hour × 3
3. After the final ethanol step, transfer the samples to a fresh 100% ethanol solution and incubate at 4 °C overnight. Note: Ensure the paraplast is melted at 58–60 °C overnight in preparation for the infiltration steps.

#### 3.2.4 Paraffin Infiltration

**Day 3–6: Infiltration (Fig. 1**)

1. Remove ethanol from the samples and replace it with Histo-Clear in a stepwise manner:

- Day 3:

- 3:1 ethanol:Histo-Clear, 1 hour
- 1:1 ethanol:Histo-Clear, 1 hour
- 1:3 ethanol:Histo-Clear, 1 hour
- 100% Histo-Clear, 1 hour × 3
- Immerse tissues in a 1:3 mixture of paraplast:Histo-Clear and incubate overnight at 60 °C.
2. Continue replacing the solution with increasing concentrations of paraplast:

- **Day 4:** 1:2, 1:1, 3:1 paraplast:Histo-Clear (3 hours each), ending with 100% paraplast overnight at 60 °C.
- **Day 5–6:** Replace with fresh 100% paraplast every 3 hours, incubating overnight at 60 °C.

Note: Always ensure sufficient melted paraplast for each infiltration step.

#### 3.2.5 Paraffin Embedding

**Day 7: Embedding (Fig. 1**)

1. Pre-warm embedding moulds on a heating plate set to 60 °C.
2. Gently stir the tissues in melted paraplast to ensure an even coating, then pour the mixture into the mould.
3. Use a needle or forceps to orient the plant tissues for sectioning (Fig. 1).
4. Allow the mould to cool on a bench or a cooling plate until the bottom solidifies, then transfer to the ice to speed up solidification.
5. Once fully hardened, remove the paraffin block from the mould and store it at 4 °C in a sealed bag.

Note: As the *Medicago* samples used were very small, 2 samples were embedded in each block to increase the opportunity to capture the correct region of interest.

### 3.3 Paraffin Sectioning and Section Slide Preparation (***12***)

**Day 8: Sectioning and section collection**

Note: all equipment should be cleaned with RNaseZap prior to sectioning and between each tissue sample.

#### 3.3.1 Tissue Block Trimming

1. Pre-chill the block on the ice tray, then mount it onto the microtome.
2. Use the microtome to trim the block to remove the outer layer of wax.

#### 3.3.2 Paraffin sectioning

1. Once the tissue has appeared at the surface of the block, take a few consecutive serial sections at 8–10 μm (note 5) (Fig. 2).
2. Using the forceps, lay these sections on the cutting mat, separate the first section from the others using the razor blade and float it onto the water bath (Fig. 2).
3. Dip a standard slide into the water bath and collect the section onto it. View this section under the microscope to check that the correct region of interest is present.
4. If the correct region is not present, discard the sections on the cutting mat and repeat the previous 3 steps, each time checking one section with the microscope (note 2).
5. Once the region of interest has been identified, return to the sections on the cutting mat, separate one section using a razor blade, and if needed, trim the surrounding wax so that the entire section fits within the Xenium capture area. If multiple sections are being collected trim the wax enough so that all sections will fit in the capture area. (Fig. 2).

Note: The region of interest in the *Medicago* tissue was very small and specific, aiming precisely for the centre of the nodule. This necessitated a cautious approach, if working with a larger region of interest, the intermediate sections may not need checking.

**Fig 2.**
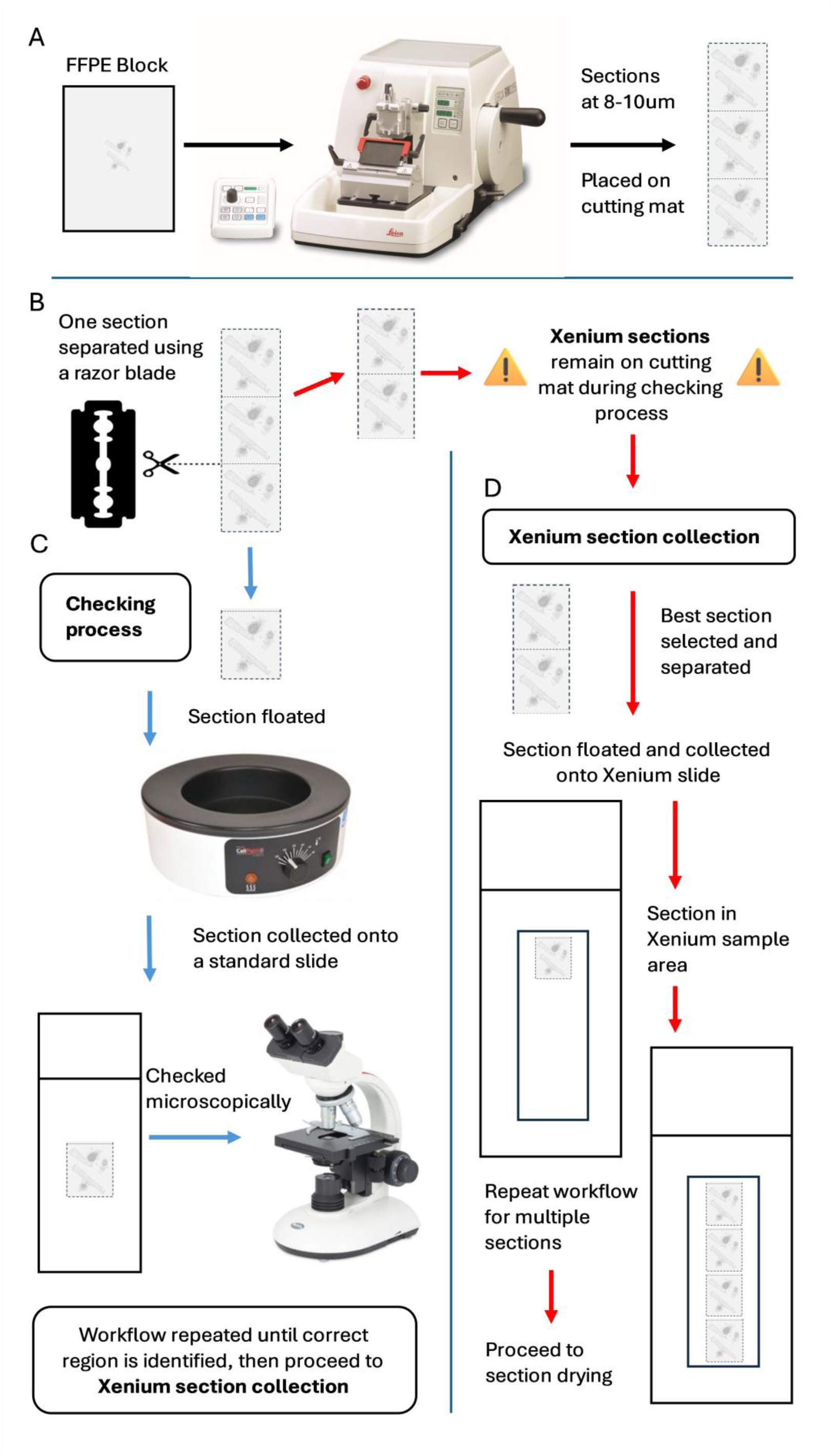
Sectioning and section collection workflow. (A) Paraffin sectioning: Use a microtome to take consecutive paraffin sections from the FFPE block, at a thickness of 8-10μm. (B) Section separation: Using a razor blade and cutting mat carefully separate one section from the set, to be used in the checking process. (C) Region of interest checking: Take the selected section and float it onto a water bath set at ∼45 °C. Dip a slide into the bath and collect the section onto it, use a microscope to check if the section shows the correct region of interest. Repeat the workflow, parts A – C until the correct region is seen. (D) Xenium section collection: Having identified the region of interest return to the remaining sections from that serial set and select the best section for Xenium. Float this section onto the water bath, dip the Xenium slide into the bath to carefully collect the section. Collect towards the top of the capture area if further sections are to be collected. Repeat the entire workflow, parts A – D, to collect additional sections onto the slide.

#### 3.3.3 Section Placement on Xenium Slides

1. Float the section on the 45 °C Milli-Q water bath for 5 – 10 seconds (note 6), then dip the Xenium slide into the bath to collect the section, ensuring to leave space for other sections if needed (Fig. 2).
2. Repeat the process for additional sections on the slide. When collecting further sections, do not dip previous sections below the waterline to keep their quality.

#### 3.3.4 Section drying and storage

1. Air dry Xenium slides for 30 minutes at room temperature in a slide rack.
2. Bake the slides for 3 hours at 42°C.
3. Transfer the slides to a closed container and dry them overnight in a desiccator at room temperature to ensure complete drying.

#### 3.3.5 Deparaffinization

1. Remove the slides from the desiccator, transfer to a slide rack, and bake in the oven for 2h at 60°C
2. Allow slide to return to room temperature, approximately 10 minutes
3. Pass the slides through a series of reagent baths to deparaffinise them:

- Xylene: 10 minutes
- Xylene: 10 minutes
- 100% ethanol: 3 minutes
- 100% ethanol: 3 minutes
- 96% ethanol: 3 minutes
- 96% ethanol: 3 minutes
- 70% ethanol: 3 minutes
- Nuclease-free water: 20 seconds

### 3.4 Probe Hybridisation and Processing Xenium Slides

#### 3.4.1 Buffer and Solution Preparation Day 1

1. 1x PBS: 10x PBS pH 7.4, Nuclease-free Water.

- Prepare a 50 mL tube of fresh 1x PBS each day by combining 45 mL of nuclease-free water and 5 mL of 10x PBS. Maintain at room temperature.
2. PBS-T: 1x PBS, 10% Tween-20, Nuclease-free Water.

- Prepare 20 mL of fresh PBS-T each day by mixing 19.9 mL of 1x PBS and 100 µL of 10% Tween-20 for the Hybridisation workflow. Make up 1 L by adding 895 mL nuclease-free water, 100 mL 10x PBS and 5 mL 10% Tween-20 for the Xenium Loading Protocol. Maintain at room temperature.
3. Diluted Perm Enzyme B: 1x PBS, Perm Enzyme B.

- Make up 1ml by using 998µl of 1x PBS and 2µl of Perm Enzyme B. Maintain at room temperature.
4. Decrosslinking Buffer: Xenium FFPE Tissue Enhancer, Urea 8M, Diluted Perm Enzyme B.

- For 2 slides + 10%, aliquot 1,017.5µl of FFPE Tissue Enhancer, then add 68.8µl of Urea 8M and top-off with 13.8µl of Diluted Perm Enzyme B. Maintain at room temperature <u>in the dark.</u>

**Day 2**:

1. Prepare fresh 1x PBS and PBS-T as above.
2. 70% Ethanol: 100% Ethanol, Nuclease-free Water.

- Make up 2ml by adding 1540µl 100% Ethanol and topping off with 660µl nuclease-free water.
3. 70% Isopropanol: 100% Isopropanol, Nuclease-free Water.

- Make up 5ml by adding 3500µl 100% Isopropanol and topping off with 1500µl nuclease-free water.
4. Xenium Probe Removal Buffer: Nuclease-free Water, DMSO 100%, KCl 2M, 10% Tween-20.

- <u>Prepare in a fume hood</u>. Using a glass bottle, add 139.5 mL of nuclease-free water, then add 150 mL of DMSO. The reaction will become warm during this step. Add 7.5ml of KCl and finally, 3ml of 10% Tween-20, mix by gentle swirling or inversion (ensure lid is secure) before transferring to the #4 Xenium Probe Removal Buffer Bottle, buffer might become warm, maintain in a fume hood at room temperature for 30 minutes to let the buffer cool down.

#### 3.4.2 Custom Probe Panel Preparation

1. Xenium Standalone Custom Probes, Xenium Add-on Custom Probes, TE Buffer pH 8.0. Centrifuge custom probe panel tubes to ensure lyophilised material is at the bottom of the tube, then resuspend according to **Table 1**.
2. Ensure the cap is firmly on after resuspension, then vortex for 30 seconds and maintain at room temperature for 5 minutes.
3. Centrifuge briefly, then prepare aliquots with at least 70 µl—sufficient for two reactions. The hybridisation step requires exactly 66 µl (for two slides), with an additional 4 µl included to ensure accurate pipetting.
4. Label the aliquots and store at -20°C.
5. For each custom standalone or add-on panel, a .json file with a unique code name is generated. This file is required for the 10X Xenium Instrument to correctly configure the run and generate the results. Ensure the Xenium Custom Probe Panel(s) .json file is uploaded to the instrument and verified before starting the run.

##### Thermal Cycler Programs

When placing the Xenium Cassettes on the Xenium Thermocycler Adaptor V1, ensure they sit securely. After closing the lid, adjust the lid until an audible click is heard. This ensures the slides remain in place on the adaptor and the lids are securely attached to the cassettes. It is advisable to set up these programs in your thermal cycler(s) prior to starting the workflow. Thermal cycling conditions are provided in **Tables 2 and 3**, with a reaction volume of 100 µl.

**Table 1.** Volumes for resuspending Custom Probes based on kit reaction size.

| Custom Probe Resuspension |  | Volume of TE Buffer, pH 8.0 (µl) |  |
| --- | --- | --- | --- |
| Kit Size |  | 4 Reaction Kit | 16 Reaction Kit |
| Xenium Add-on |  | 140 | 700 |
| Xenium Standalone |  | 140 | 700 |

**Table 2.** Thermal cycler programs for the first 3 protocol steps: Decrosslinking, Probe Hybridisation and Post-Hybridisation Wash.

| Protocol | 1. Decrosslinking | 2. Probe Hyb | 3. Post-Hyb Wash |
| --- | --- | --- | --- |
| Lid Temp | 80°C | 50°C | 37°C or Off |
| Step |  |  |  |
| 1. | 22°C – Hold | 50°C – Hold | 37°C – Hold |
| 2. | 80°C – 00:30:00 | 50°C – 16:00:00 | 37°C – 00:30:00 |
| 3. | 22°C – 00:10:00 | 50°C – 08:00:00 | 37°C – Hold |
| 4. | 22°C – Hold | 50°C – Hold | N/A |

**Table 3.**
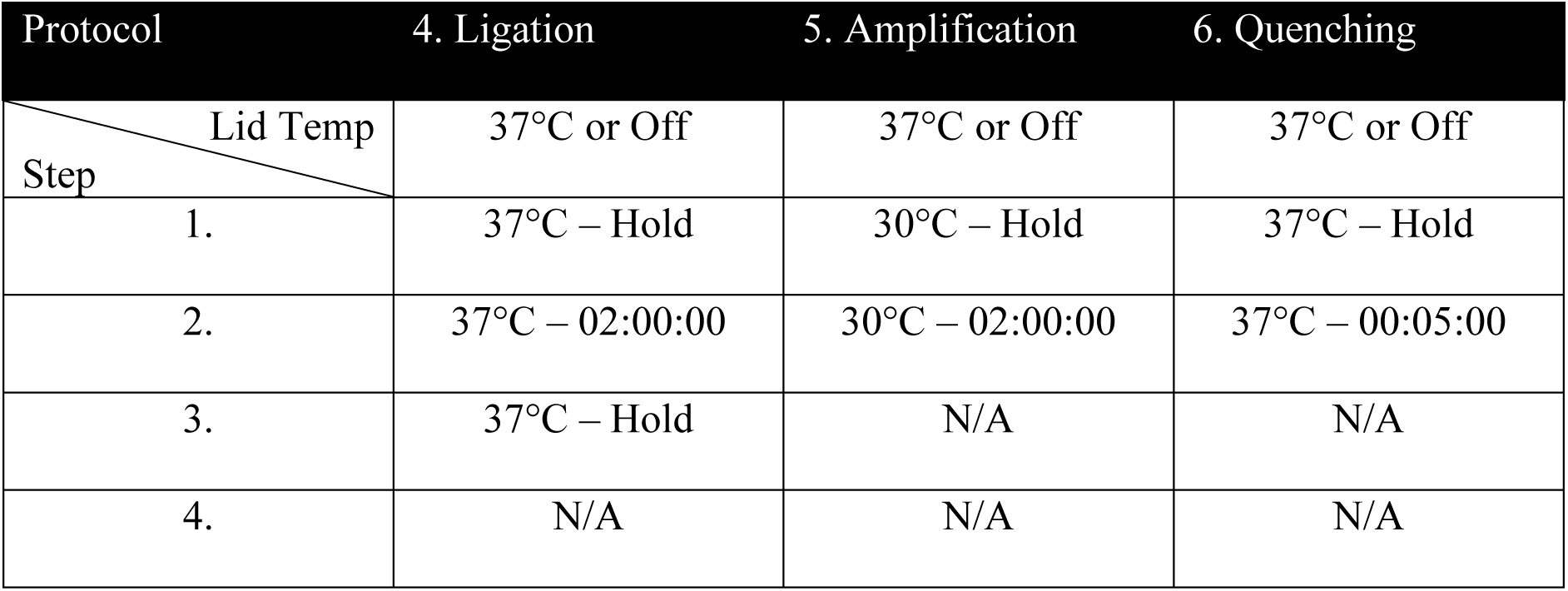
Thermal cycler programs for the last 3 protocol steps: Ligation, Amplification and Autofluorescence Quenching.

| Protocol | 4. Ligation | 5. Amplification | 6. Quenching |
| --- | --- | --- | --- |
| Lid Temp | 37°C or Off | 37°C or Off | 37°C or Off |
| Step |  |  |  |
| 1. | 37°C – Hold | 30°C – Hold | 37°C – Hold |
| 2. | 37°C – 02:00:00 | 30°C – 02:00:00 | 37°C – 00:05:00 |
| 3. | 37°C – Hold | N/A | N/A |
| 4. | N/A | N/A | N/A |

#### 3.4.3 Decrosslinking

Before starting, clean all surfaces, micropipettes, and pipette controllers with RNase AWAY and ensure all glassware used is autoclaved. Ensure you are wearing the correct personal protective equipment (PPE), including a lab coat and gloves. All volumes listed in these methods are prepared for two Xenium Slides, but more can be prepared and stored according to sections 3.4.9 and 3.4.10.

Decrosslinking is a crucial step when working with formalin-fixed embedded material, enabling accurate probe hybridisation by reversing the covalent bonds formed between DNA, RNA, and proteins during the fixation step. When slides are undergoing the deparaffinization process, prepare all buffers for Decrosslinking and Probe Hybridisation steps, 1x PBS, PBS-T, Decrosslinking Buffer and Xenium Cassettes.

1. Remove the slides from the glass Coplin jars containing nuclease-free water and wipe the back of the slides with lint-free laboratory wipes to remove any remaining water drops. Place slides in the Xenium Cassettes with the top clicking into place, then gently aliquot 500µl 1x PBS into the corner of the sample area without touching or damaging the tissue section(s).
2. Place the Xenium Thermocycler Adaptor V1 in the thermal cycler and start the decrosslinking protocol listed in **Table 4** to equilibrate to the required temperature:
3. Remove the 1x PBS from the slides and add 500 µL of Decrosslinking Buffer, avoiding bubbles. Gently tap the slides on the lab bench to ensure even coverage.
4. Place a Xenium Cassette Lid on the Xenium Cassette, place the cassette on the thermal cycler at 22°C and skip the “Hold” step to begin the Decrosslinking.
5. During this incubation, retrieve the Probe Hybridisation reagents and follow the preparation and handling steps for each:

a. Probe Hybridisation Buffer: Thaw at room temperature for ∼15 minutes, then check for precipitate and invert till clear. Maintain at room temperature.
b. Xenium Standalone and Add-on Probe Resuspended Aliquots: Thaw and maintain at room temperature.
c. Preheat a heat block to 95°C.
6. Once the Decrosslinking Protocol is finished, retrieve the Xenium Cassettes from the thermal cycler.
7. Remove the Xenium Cassette Lids and remove all Decrosslinking Buffer from the sample area. Discard the lids after removal. Check the tissue for any signs of detachment—if detachment is observed, stop the workflow and contact 10x Genomics Support for immediate guidance.
8. Add 500 µL of PBS-T to the well and incubate for 1 minute at room temperature. Then, remove all PBS-T.
9. Repeat step 8 twice for a total of 3 washes.
10. Add 500µl of PBS-T to the well.

**Table 4.** The Decrosslinking thermal cycler program with details of each step.

| Lid Temperature | Reaction Volume | Run Time |
| --- | --- | --- |
| 80°C | 100µl | 40 Minutes |
| **Step** | **Temperature** | **Time** |
| Hold | 22°C | Hold |
| Decrosslinking | 80°C | 00:30:00 |
| Re-equilibrate | 22°C | 00:10:00 |
| Hold | 22°C | Hold |

#### 3.4.4 Probe Hybridisation

Xenium employs the use of padlock-type probes consisting of two 20-base RNA-binding arms connected by a backbone. When both arms successfully hybridize to the target RNA, ligation occurs, forming a circular template for rolling circle amplification and generating a unique barcode (Fig. 3).

1. Place an aliquot of each of the probe panels in the heat block at 95°C for 2 minutes, immediately after remove to ice for 1 minute, then maintain on ice.
2. Prepare the Probe Hybridisation Mix as per **Table 5** by adding in the order listed, pipette mix 15 times, and centrifuge briefly, then maintain at room temperature.
3. Prepare a thermal cycler with the Probe Hybridisation Protocol listed in **Table 6** and start the Pre-equilibrate step.
4. Remove all the PBS-T, aspirate remaining buffer using a P20 tilting the cassette to ensure all buffer is removed.
5. Add 500µl of the Probe Hybridisation Mix to the sample area without introducing bubbles and gently tap the cassette on the bench to even the coverage.
6. Apply a new Xenium Cassette Lid, then place it on the pre-equilibrated thermal cycler.
7. Skip the Pre-equilibrate step to initiate the overnight Probe Hybridisation.

**Fig. 3.**
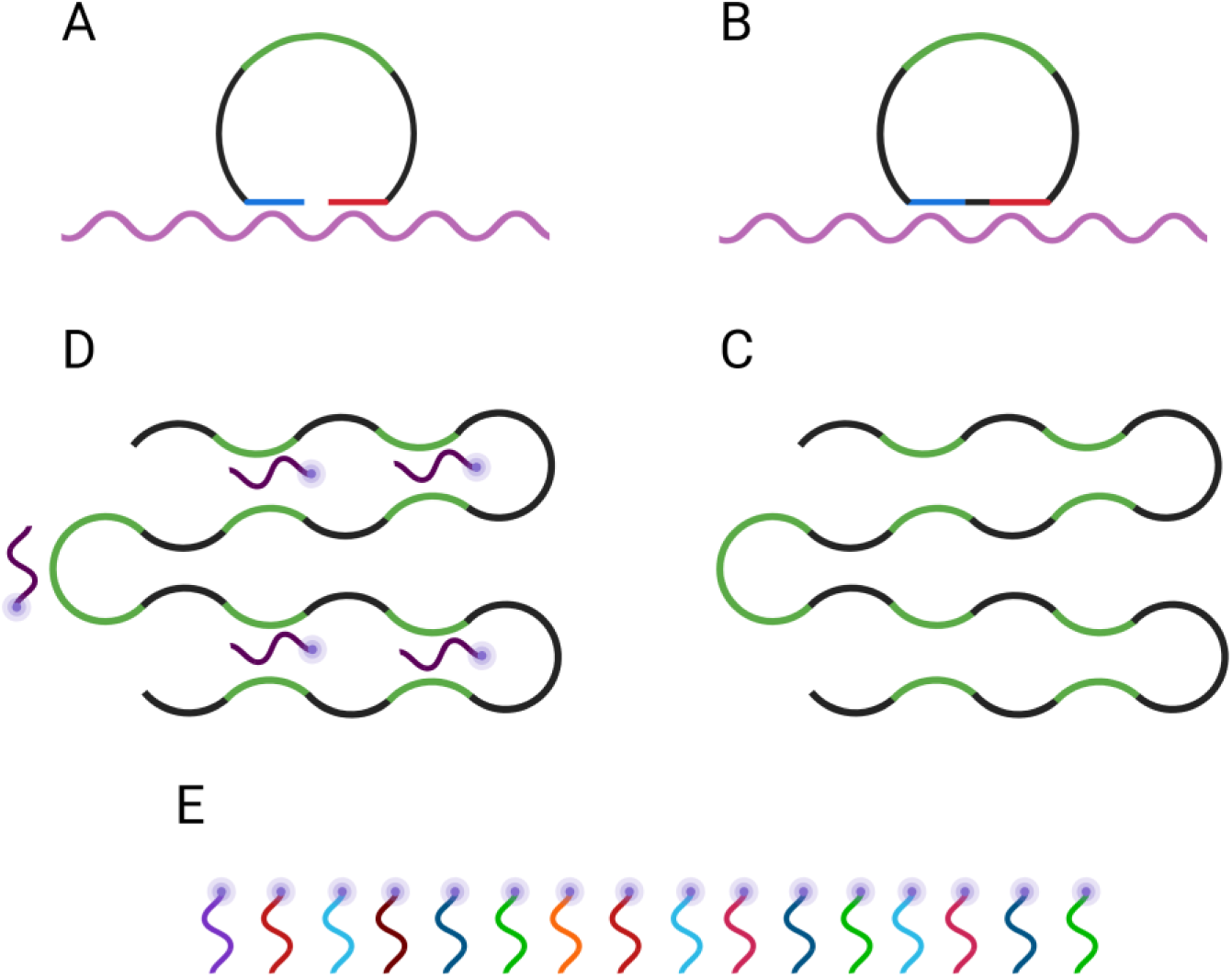
Workflow for hybridisation, amplification, and detection of RNA targets using Xenium in situ. (A) A padlock probe hybridises to the RNA target, with left and right arms (red and blue) complementary to adjacent regions on the transcript. The top region (green) is a target-specific sequence. (B) Ligation of the probe generates a circular DNA molecule. (C) Rolling circle amplification of the circularised probe produces a concatemer containing multiple repeats of the target-specific sequence. (D) Fluorescently labelled detection oligonucleotides are introduced within the Xenium Analyser to hybridise to the amplified target sequence. (E) Sequential cycles of oligonucleotide hybridisation and imaging reveal a unique barcode (“codeword”) corresponding to each RNA target (*14*).

**Table 5.** The reagents and volumes required for 2 reactions of the Probe Hybridisation Mix.

| **Probe Hybridisation Mix** | **Volume for 2 Reactions + 5% (µl)** |
| --- | --- |
| Probe Hybridisation Buffer | 630 |
| TE Buffer, pH 8.0 | 288 |
| Xenium Standalone Custom Probes | 66 |
| Xenium Add-on Custom Probes | 66 |
| **Total** | 1050 |

**Table 6.** The Probe Hybridisation thermal cycler program with details of each step. A minimum of 16 hours is required for effective probe hybridisation.

| **Lid Temperature** | **Reaction Volume** | **Run Time** |
| --- | --- | --- |
| 50°C | 100µl | 16-24 Hours |
| **Step** | **Temperature** | **Time** |
| Pre-equilibrate | 50°C | Hold |
| Probe Hybridisation I | 50°C | 16:00:00 |
| Probe Hybridisation II | 50°C | 08:00:00 |
| Hold | 50°C | Hold |

Note: If planning to load the Xenium Slides onto the Xenium Analyser the next day, retrieve the Decoding Reagent Module B from the -20°C storage and place it at 4°C overnight.

#### 3.4.5 Post-Hybridisation Wash

The post-hybridisation wash uses a gentle detergent to remove excess probes and minimise background noise.

1. Retrieve the Post-Hybridisation Wash Buffer and thaw at room temperature for 30 minutes. Vortex and centrifuge briefly. Maintain at room temperature.
2. Prepare fresh 1x PBS and PBS-T; do not use any leftovers from the previous day.
3. For the next wash steps, remove and add buffers quickly. When working with 2 slides, remove and add buffers to 1 slide at a time.
4. Open the thermal cycler lid and check for slide cracks or leaked fluid. If there are any signs of leakage or glass cracks, contact 10x Genomics support immediately.
5. Retrieve the Xenium Cassettes and remove and discard the Cassette Lids.
6. Remove all Probe Hybridisation Mix from the corner of the sample area and immediately add 500 µL PBS-T. Incubate for 1 minute at room temperature.
7. Remove all PBS-T and add 500µl PBS-T, incubate for 1 minute at room temperature, and while incubating, prepare a thermal cycler with the following incubation protocol and start the Pre-equilibrate step (Table 7).
8. Remove all PBS-T and add 500µl to the sample area without introducing bubbles.
9. Apply a new Xenium Cassette Lid on each cassette and on the pre-heated thermal cycler before closing the lid and skipping the Pre-equilibrate step.
10. During this incubation, retrieve the Probe Ligation reagents and follow the preparation and handling steps for each:

a. Probe Ligation Buffer: Thaw at room temperature for ∼15 minutes, pipette mix and centrifuge briefly. Maintain at room temperature.
b. Ligation Enzyme A: Pipette mix and centrifuge briefly. Maintain on ice. Ligation Enzyme B: Pipette mix and centrifuge briefly. Maintain on ice.

**Table 7.** The Post-Hybridisation thermal cycler program with details of each step.

| **Lid Temperature** | **Reaction Volume** | **Run Time** |
| --- | --- | --- |
| 37°C or Off | 100µl | 30 Minutes |
| **Step** | **Temperature** | **Time** |
| Pre-equilibrate | 37°C | Hold |
| Post-Hybridisation Wash | 37°C | 00:30:00 |
| Hold | 37°C | Hold |

#### 3.4.6 Ligation

Once the padlock probes successfully anneal to the RNA target sequence, ligation occurs, creating a circular product that serves as a template for subsequent amplification.

1. Prepare the Ligation Mix by adding the reagents in the order listed in **Table 8**. Then, pipette the mix 10 times and centrifuge briefly. Maintain on ice.
2. Remove Xenium Cassette(s) from thermal cycler and remove the Xenium Cassette Lids.
3. Remove all Post-Hybridisation Wash Buffer and discard the Lids.
4. Immediately add 500µl PBS-T and incubate for 1 minute at room temperature. Remove all PBS-T.
5. Repeat step 4 once more.
6. Add 500μl PBS-T. Incubate for 1 min at room temperature.
7. During final incubation, prepare a thermal cycler with the Ligation protocol listed in **Table 9** and start the Pre-equilibrate step.
8. Remove all PBS-T from the well, using a P20 pipette to aspirate any remaining liquid from the corners of the well.
9. Add 500μl Ligation Mix to each well and apply a new Xenium Cassette Lid.
10. Place cassette(s) on pre-heated thermal cycler, close the lid and skip the Pre-equilibrate step to initiate the protocol.

**Table 8.** The reagents and volumes required for 2 reactions of the Ligation Mix.

| Ligation Mix | Volume for 2 Reactions + 10% (μl) |
| --- | --- |
| Ligation Buffer | 962.5 |
| Ligation Enzyme A | 27.5 |
| Ligation Enzyme B | 110 |
| **Total** | **1100** |

**Table 9.**
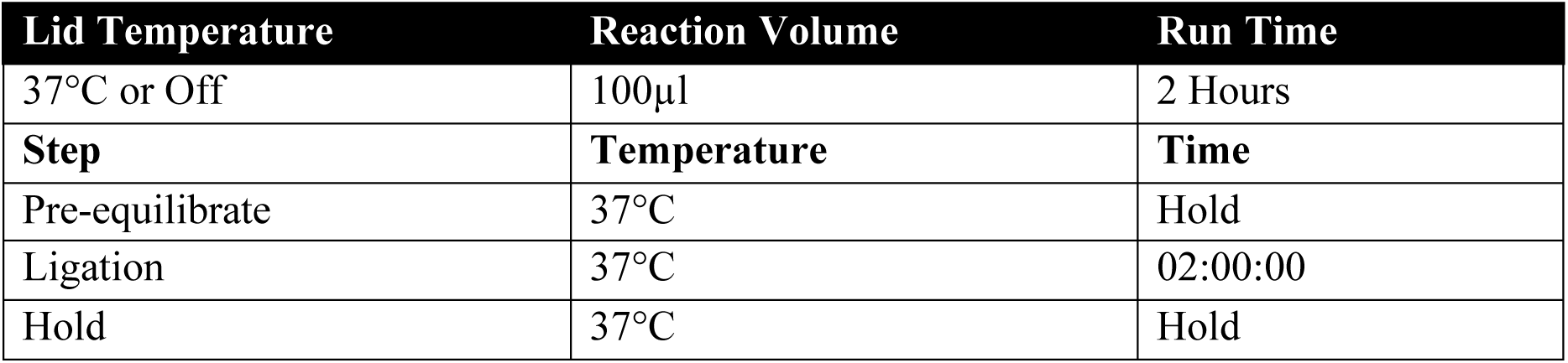
The Ligation thermal cycler program with details of each step.

| Lid Temperature | Reaction Volume | Run Time |
| --- | --- | --- |
| 37°C or Off | 100μl | 2 Hours |
| Step | Temperature | Time |
| Pre-equilibrate | 37°C | Hold |
| Ligation | 37°C | 02:00:00 |
| Hold | 37°C | Hold |

Once the incubation begins, retrieve the Amplification Mix and thaw it on ice. Once thawed, vortex then centrifuge and return to ice.

#### 3.4.7 Amplification

The circular template undergoes rolling circle amplification to enhance the signal for detection and decoding on the Xenium Analyzer, ensuring that multiple copies of the barcode are generated and resolved.

1. Retrieve the Amplification Enzyme immediately before use. Pipette the mix, then centrifuge briefly and maintain on ice.
2. Prepare the Amplification Master Mix by adding the reagents in the order listed in **Table 10**. Then, pipette the mix 10 times and centrifuge briefly. Maintain on ice:
3. Remove Xenium Cassette(s) from thermal cycler and remove the Xenium Cassette Lids.
4. Remove all Ligation Mix and discard the Lids.
5. Immediately add 500µl PBS-T and incubate for 1 minute at room temperature. Remove all PBS-T.
6. Repeat step 5 once more.
7. Add 500μl PBS-T. Incubate for 1 min at room temperature.
8. During final incubation, prepare a thermal cycler with the Amplification protocol listed in **Table 11** and start the Pre-equilibrate step:
9. Remove all PBS-T, and use a P20 to remove any remaining PBS-T.
10. Add 500µl Amplification Master Mix to the well and apply a new Xenium Cassette Lid.
11. Place cassette(s) on pre-heated thermal cycler, close the lid and skip the Pre-equilibrate step to initiate the protocol.
12. With 20 minutes remaining on this incubation, retrieve the Autofluorescence Mix and begin thawing in a thermomixer set to 37°C and 300 revolutions per minute (rpm) for 15 minutes, then cool to room temperature for 5 minutes. Once thawed, vortex and then centrifuge, maintaining at room temperature.
13. Retrieve Reducing Agent B and (if completing protocol up to section 3.4.10 on the same day) Nuclei Staining Buffer, thaw both at room temperature, then vortex and centrifuge briefly. Maintain at room temperature.

**Table 10.** The reagents and volumes required for 2 reactions of the Amplification Master Mix.

| Amplification Master Mix | Volume for 2 Reactions + 10% (μl) |
| --- | --- |
| Amplification Mix | 990 |
| Amplification Enzyme | 110 |
| **Total** | **1100** |

**Table 11.** The Amplification thermal cycler program with details of each step.

| Lid Temperature | Reaction Volume | Run Time |
| --- | --- | --- |
| 37°C or Off | 100μl | 2 Hours |
| Step | Temperature | Time |
| Pre-equilibrate | 30°C | Hold |
| Amplification | 30°C | 02:00:00 |

#### 3.4.8 Post Amplification Wash

The additional wash removes the amplification master mix to prevent interaction with the subsequent quenching.

1. Remove Xenium Cassette(s) from thermal cycler and remove the Xenium Cassette Lids.
2. Remove all Amplification Mix and discard the Lids.
3. Immediately add 500µl TE Buffer pH 8.0 and incubate for 1 minute at room temperature. Remove all TE Buffer.
4. Repeat step 3 once.
5. Add 500µl TE Buffer pH 8.0. Incubate for 1 min at room temperature.

#### 3.4.9 Autofluorescence Quenching

Autofluorescence refers to background signals generated by the tissue and/or sample processing. Autofluorescence quenching reduces these unwanted signals and enhances the signal-to-noise ratio, ensuring the generation of high-quality data.

1. Prepare the solutions in **Table 12**.
2. Remove all TE Buffer.
3. Immediately add 1,000µl PBS-T and incubate for 1 minute at room temperature. Remove all PBS-T.
4. Repeat step 3 twice for a total of 3 washes
5. Add 500µl Diluted Reducing Agent B and apply a new Xenium Cassette Lid, then incubate for 10 minutes at room temperature.
6. Remove but keep the Xenium Cassette Lid and remove all Diluted Reducing Agent B.
7. Add 1000µl **<u>70% Ethanol</u>** and incubate for 1 minute at room temperature. Remove all 70% Ethanol.
8. Add 1000µl **<u>100% Ethanol</u>** and incubate for 1 minute at room temperature. Remove all 100% Ethanol.
9. Repeat step 8 once more.
10. Pipette mix the Autofluorescence Solution 5 times, then add 500 µL to each well and incubate for 10 minutes at room temperature <u>in the dark.</u>
11. During the incubation, prepare a thermal cycler with the following Drying protocol (**Table 13**) and start the pre-equilibration.
12. Remove and discard the Xenium Cassette Lids, then remove all Autofluorescence Solution.
13. Add 1000µl 100% Ethanol and incubate for 2 minutes at room temperature. Remove all 100% Ethanol.
14. Repeat step 13 twice more.
15. Place the Xenium Cassette without a lid on the pre-equilibrated thermal cycler and skip the step to initiate the Drying protocol.
16. Immediately remove the Xenium Cassette once the protocol has finished and add 1000µl 1X PBS, then incubate for 1 minute at room temperature <u>in the dark.</u>
17. Remove all 1x PBS and add 1000µl <u>PBS-T,</u> then incubate for 2 minutes at room temperature <u>in the dark.</u>

**Table 12.** The reagents and volumes required for 2 reactions of the 3 Autofluorescence Quenching Solutions. A. Diluted Reducing Agent B: Add reagents in the listed order, then vortex and maintain them at room temperature. B. 70% Ethanol, add reagents in the listed order, then vortex and maintain at room temperature. C. Autofluorescence Solution: Add reagents in the listed order, then vortex and maintain at room temperature <u>in the dark</u>.

| A. Diluted Reducing Agent B |  | B. 70% Ethanol |  | C. Autofluorescence Solution |  |
| --- | --- | --- | --- | --- | --- |
| Reagent | Volume (µl) | Reagent | Volume (µl) | Reagent | Volume (µl) |
| 1x PBS | 1089 | NFW | 660 | 100% Ethanol | 1089 |
| Reducing Agent B | 11 | 100% Ethanol | 1540 | Autofluorescence Mix | 11 |
| **Total** | **1100** | **Total** | **2000** | **Total** | **1100** |

**Table 13.**
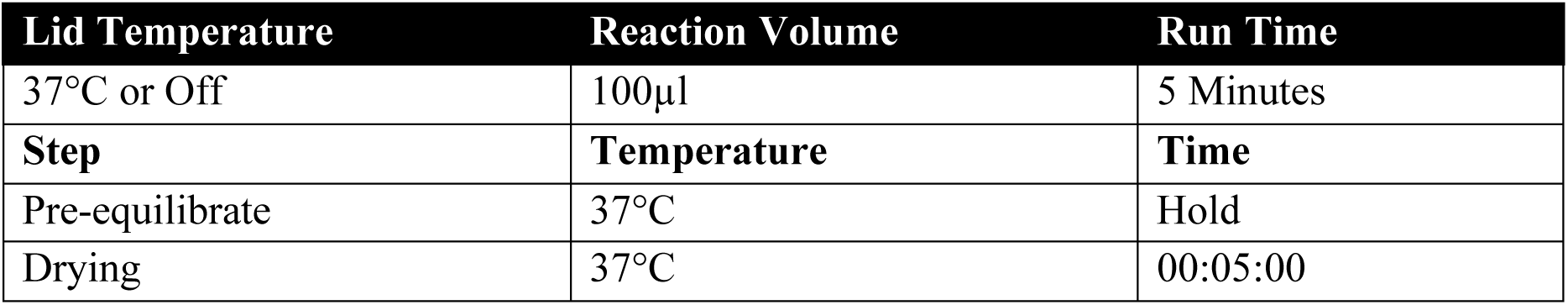
The Autofluorescence Quenching thermal cycler program with details of each step.

| Lid Temperature | Reaction Volume | Run Time |
| --- | --- | --- |
| 37°C or Off | 100µl | 5 Minutes |
| Step | Temperature | Time |
| Pre-equilibrate | 37°C | Hold |
| Drying | 37°C | 00:05:00 |

Safe Stop: Slides can be stored at 4°C in the dark for 16-24 hours. Alternatively, continue immediately.

#### 3.4.10 Nuclei Staining

DAPI staining is a key component used to stain nuclei, aiding in both tissue or region of interest identification and cell segmentation.

1. If stored, remove the Xenium Cassette Lid. Remove all PBS-T from the well, save the lid for use in the following indicated steps.
2. Add 500μl Nuclei Staining Buffer and incubate 1 minute at room temperature <u>in the dark</u>.
3. Remove Nuclei Staining Buffer.
4. Add 1000µl PBS-T, then incubate for 1 minute at room temperature <u>in the dark</u>. Remove all PBS-T.
5. Repeat step 4 twice more.
6. Add 1000µl PBS-T.
7. Proceed to 3.4.11 and 3.4.12 Reagent Preparation and Loading the Xenium or, at this point, slides can be stored for up to one week at 4°C in the dark with a new Xenium Cassette Lid and replacement of PBS-T every 3 days, or for up to 2 months at -20°C after performing the following dehydration and cryoprotection steps:
8. Prepare 10 mL 30% Glycerol in PBS
9. Remove all PBS-T.
10. Add 1000µl 70% Ethanol and incubate for 2 minutes at room temperature. Remove all 70% Ethanol.
11. Repeat step c twice more.
12. Remove slides from Xenium Cassette and place in a slide mailer containing 10ml 30% Glycerol in PBS.
13. When ready to use, equilibrate the slide to room temperature for 30 minutes.
14. Rinse mailer with 10 mL PBS-T three times.
15. Remove slide from mailer and place in Xenium Cassette, add 1000µl PBS-T.

#### 3.4.11 Xenium V1 Workflow Reagent Preparation – Imaging and Decoding

The Xenium Analyzer uses a high-resolution imaging system to detect amplified probes. During the imaging and decoding step, multiple rounds of fluorescent probe hybridisation, imaging, and probe removal are carried out to generate a unique optical barcode for each gene. The Xenium Analyzer run requires several specialised buffers to ensure accurate and efficient in situ sequencing (ISS).

1. If Decoding Reagent Module B was not thawed overnight at 4 °C, place it in a water bath at 37°C for 2.5 hours.
2. If Module B is already thawed, proceed to step 3 immediately. If thawing is required, use the final 45 minutes of incubation to prepare the following buffers.
3. Add 500 µl of Nuclease-free Water to Decoding Reagent Bottle 1 (the “green bottle”) provided in the kit.
4. In a 1 L glass bottle, prepare 1x PBS-T as described in **Table 14** and transfer it to Decoding Reagent Bottle 2 (the “yellow bottle”).
5. Add 150 µl of Nuclease-free Water to Decoding Reagent Bottle 3 (the “red bottle”).
6. In a fume hood, prepare the Xenium Probe Removal Buffer using a glass bottle, following the order of ingredients listed in **Table 15**. Cap and gently invert the bottle 10 times to mix. Let the solution cool at room temperature for 30 minutes, then transfer it to Decoding Reagent Bottle 4 (the “blue bottle”) and proceed with Xenium loading.
7. After the 30-minute incubation, replace all Decoding Reagent Bottle lids with the Xenium Buffer Caps.
8. Start a Xenium run on the Xenium Analyser, which will initiate a ∼5-minute self-check.
9. Retrieve both the Decoding Reagent Modules A and B from 4°C storage, or Decoding Module B from the water bath if not thawed overnight.
10. Remove the mylar packaging and remove both lids and elastic bands, then mix by inverting 20 times.
11. Centrifuge Decoding Reagent Module A at 1600 relative centrifugal force (rcf) for 1 minute at room temperature, then place on ice.
12. Centrifuge Decoding Reagent Module B at 300 rcf for 1 minute at room temperature, then maintain at room temperature.

**Table 14.**
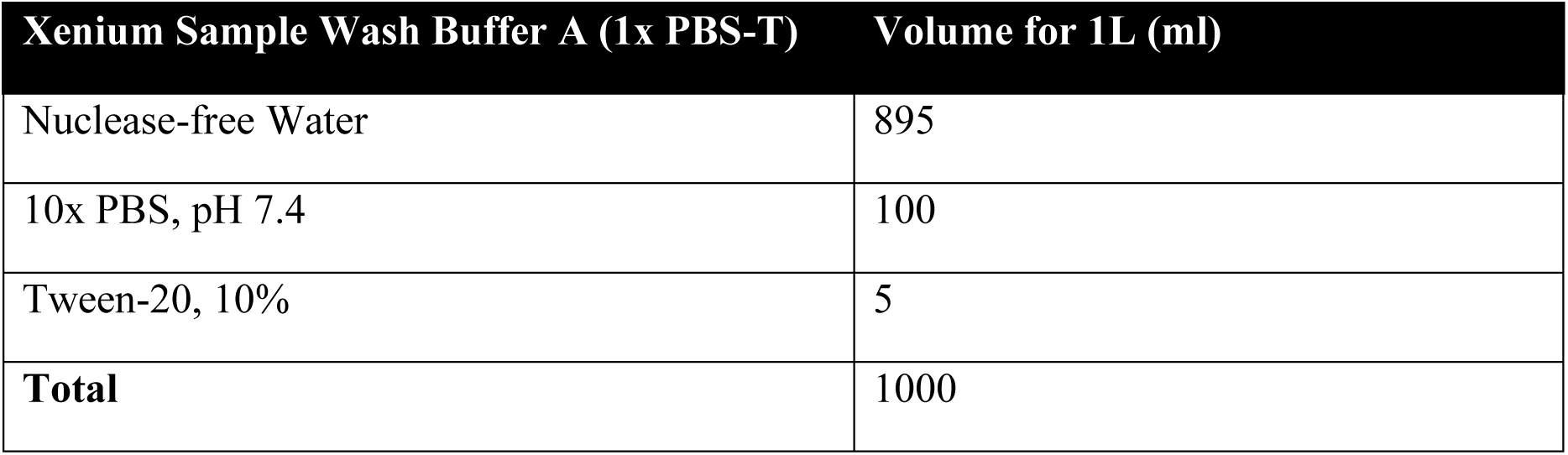
The reagents and volumes required for the Xenium Sample Wash Buffer A.

| Xenium Sample Wash Buffer A (1x PBS-T) | Volume for 1L (ml) |
| --- | --- |
| Nuclease-free Water | 895 |
| 10x PBS, pH 7.4 | 100 |
| Tween-20, 10% | 5 |
| **Total** | **1000** |

**Table 15.** The reagents and volumes required for the Xenium Probe Removal Buffer.

| Xenium Probe Removal Buffer | Volume (ml) |
| --- | --- |
| Nuclease-free Water | 139.5 |
| DMSO | 150 |
| KCL 2M | 7.5 |
| Tween-20, 10% | 3 |
| **Total** | 300 |

#### 3.4.12 Loading the Xenium Analyser and Initiating a Run

1. Retrieve the Decoding Consumables.
2. Once the Xenium self-check is complete, fill in the necessary run and Cassette information:

a. Run Name.
b. Select V1 or Prime workflow.
c. Indicate whether the Cell Segmentation Kit is used (not applicable for this workflow)
d. Select the on-board analysis version
e. Enter Xenium Cassette details, including Sample Name and Xenium Slide ID.
f. Specify the panel used—either Pre-loaded or Custom.
3. The Xenium Analyser door will unlock and provide instructions on loading the reagents, with more detailed instructions and a visual aid if selecting a specific consumable (Fig. 4).
4. Place each of the Decoding Reagent Bottles (1-4) in the appropriate slots. It is imperative that the Decoding Reagent Bottles are placed in the correct positions and the Xenium Buffer Caps are engaged by lowering the stoppers and securing them to the Buffer Caps (Fig. 4).
5. Slot the Decoding Modules A and B into their position, ensuring the orientation is correct and they sit <u>flat</u>.
6. Unwrap and place the Pipette Tip Rack into its slot until you feel a click, ensuring the lid is removed.
7. Place the Extraction Tip on the Liquid Handling Arm with a gentle twist, then place the Objective Wetting Consumable below it in the correct orientation.
8. Ensure that there are no tips from previous runs in the Waste Tip Tray and that it is present and properly slotted.
9. Spray the Xenium Cassette Stage with compressed air to clean.
10. Use 70% isopropanol to clean the Cassette Stage.
11. Clean the underside of the Xenium Slides with 70% isopropanol. Allow the slides to dry for a few seconds, and ensure they are free of residue or debris before proceeding.
12. Place the Xenium Slides in the correct location as dictated by your previous choice during the run set-up, such that the Slide IDs match and ensure they sit comfortably and are without cracks or leaks.
13. Remove and store the Xenium Cassette Lids, then close the Xenium Cassette Stage Lid to lock the Slides in place.
14. Use the screen to confirm all steps are complete, then shut the door.
15. Press Continue, which will initiate a ∼40-minute scan.
16. After the scan is complete, use the touchscreen interface to select regions of interest (ROIs). You may assign names or IDs to each region.
17. There are a few available slider tools to change the intensity of the Nuclei and Autofluorescence stains to help identify regions or better define morphology.
18. Ensure there are no overlapping regions and that all relevant tissue is captured. If tissue extends to the edge of the field of view, make sure all corresponding fields are selected.
19. Once regions have been selected for both cassettes, press **Continue**.
20. Review and verify all run settings and cassette details before proceeding.
21. Clicking “Start Run” will initiate the Xenium Run and provide a fairly accurate estimated time of completion.
22. Once the run is complete, select **“Start Cleanup”**. All output files will be generated per region, including a summary QC HTML report, and will be ready for downstream analysis.
23. After cleanup is finished, the instrument door will unlock, and reagents and consumables can be safely disposed of.
24. Place the Xenium Slides on a flat surface, remove any remaining liquid from the surface, then add 1 ml of PBS-T. Cover the slides with the cassette lids saved from step 13. Slides can be stored at 4 °C for up to one week.

### 3.5 Post-Xenium Processing

Because tissue integrity is preserved during and after the Xenium run, common post-Xenium workflows such as immunofluorescence (IF) staining and hematoxylin and eosin (H&E) staining can be applied. In this protocol, we used H&E staining for histological analysis and confocal microscopy to capture intrinsic cell wall autofluorescence, which aids in cell segmentation.

**Fig. 4.**
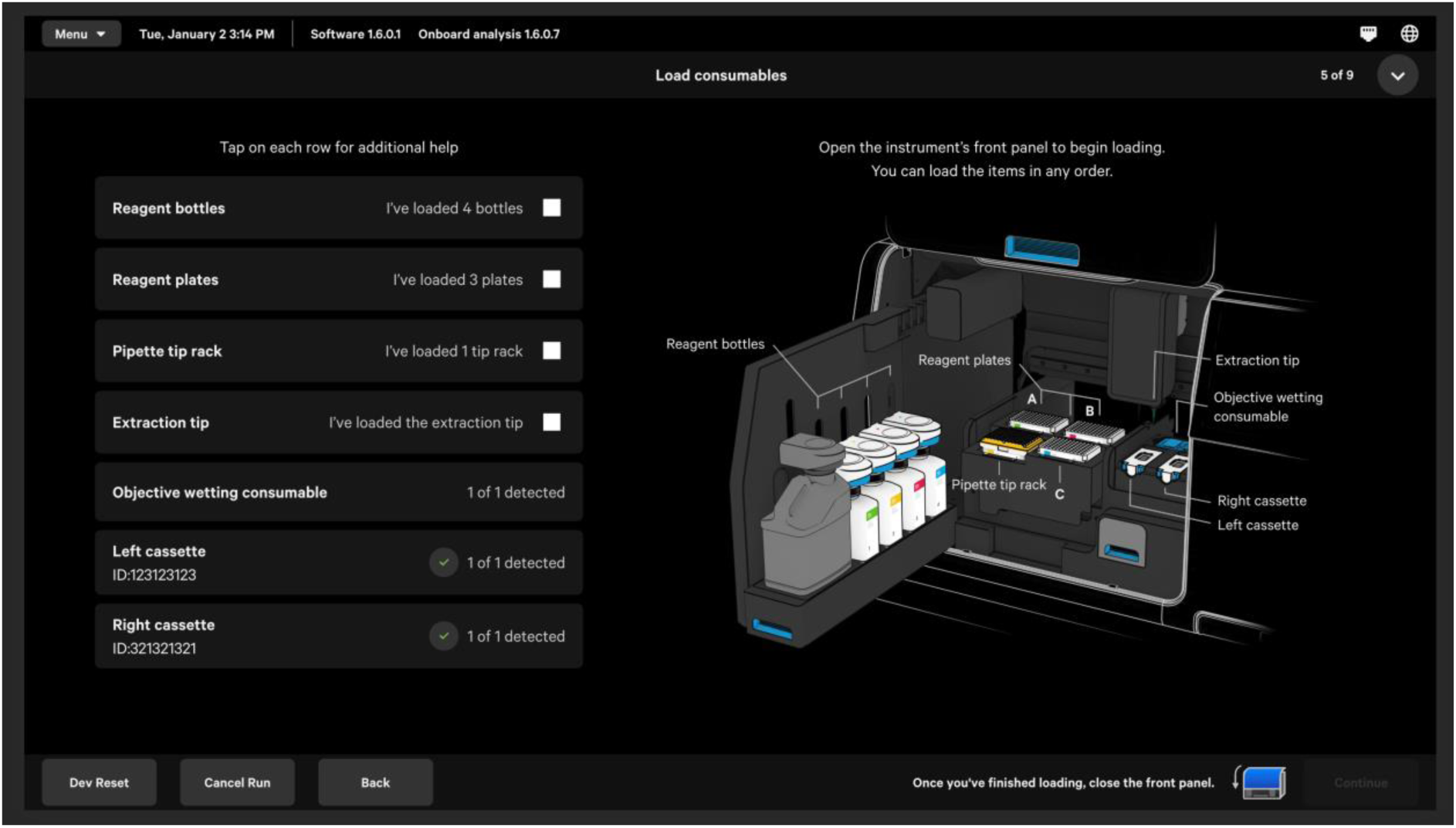
Loading interface of the Xenium Analyser. The interface displays a checklist of required reagents, each visualised in its designated position on the instrument. Selecting a checklist item reveals additional loading instructions. The first four rows require manual confirmation by the user, while the remaining three are automatically verified by the Xenium Analyser through onboard detection (*15*).

#### 3.5.1 Post-Xenium H&E Staining Quencher removal

1. Prepare the quencher removal solution:

1. Weigh 17.4 mg of Sodium Hydrosulfite and add to 10 mL of Milli-Q water
2. Close the container and vortex for 10 seconds
2. Remove the slide from the Xenium cassette and pass through a series of reagent baths:

1. Quencher removal solution: 10 minutes
2. Milli-Q water: 1 minute
3. Milli-Q water: 1 minute
4. Milli-Q water: 1 minute

**H&E staining** (Fig. 5C)

1. Load the slide onto the automated stainer, which will pass the slide through a series of pre-programmed reagent baths:

1. Harris haematoxylin: 2 minutes
2. Running tap water: 5 minutes
3. 2% Acid alcohol: 20 seconds
4. Running tap water: 5 minutes
5. Eosin: 7 minutes
6. Running tap water: 15 seconds
7. Ethanol 50% + 0.1% Tween 20: 20 seconds
8. Ethanol 70% + 0.1% Tween 20: 20 seconds
9. Ethanol 100%: 30 seconds
10. Ethanol 100%: 1 minute
11. Xylene: 5 minutes
12. Xylene: 5 minutes
2. Load the slide onto the automated coverslipper, which will dispense DPX mountant onto the slide and place a coverslip on top.

**Fig. 5.**
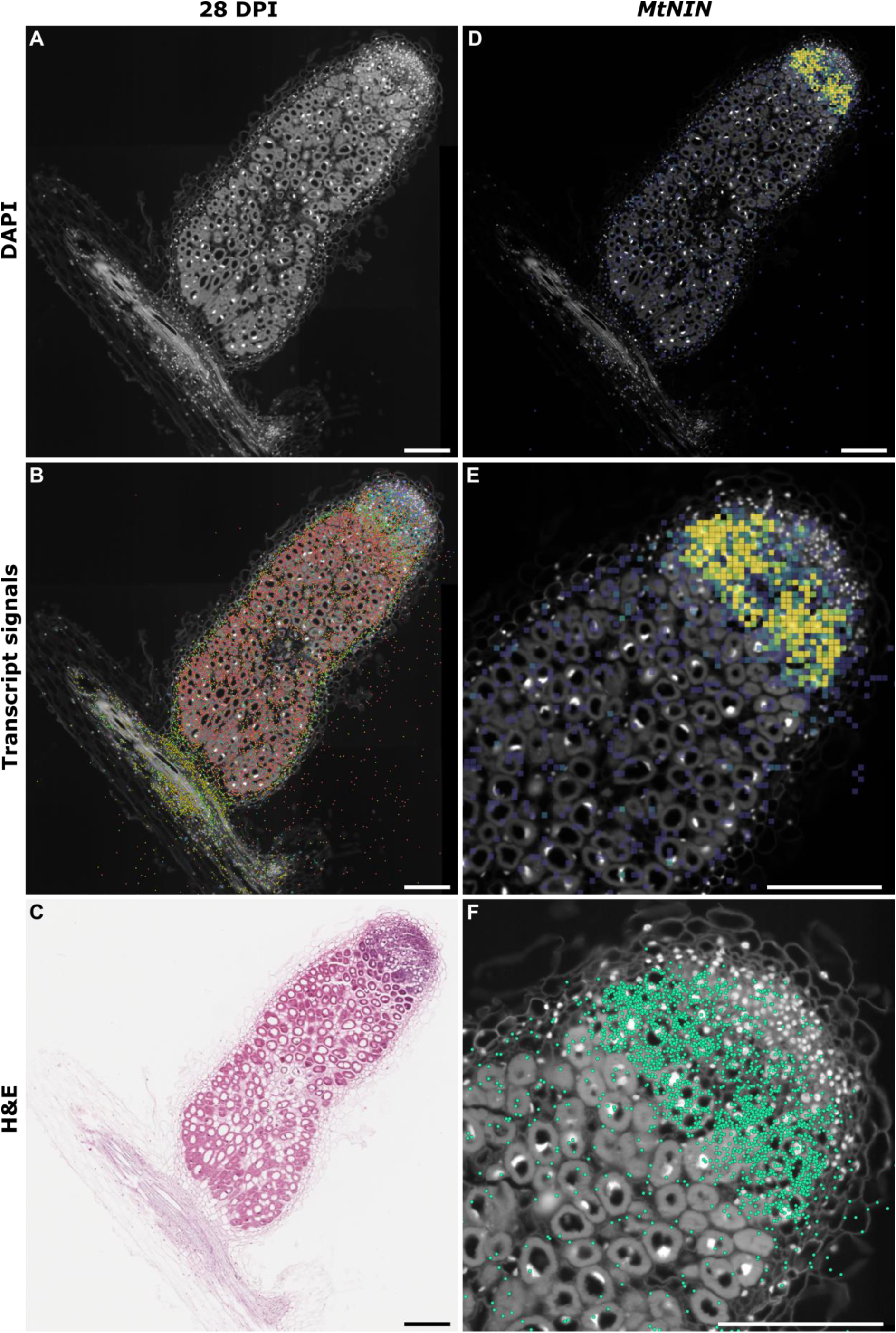
Xenium in situ sequencing performance in *Medicago* mature nodules. (A–C) An 8 μm FFPE section from a 28-day post-inoculation *Medicago* nodule. (A) A Z-slice view at the midpoint of a 10-step Z-stack shows cell wall autofluorescence and DAPI-stained nuclei. (B) Expression patterns of 50 selected genes were used to identify major cell types. Different colours represent individual genes. (C) Hematoxylin and eosin (H&E) staining performed post-Xenium workflow highlights preserved tissue integrity. (D-E) *Medicago NODULE INCEPTION (MtNIN)* expression density maps with a bin size of 10 μm. The colour scale ranges from purple (lowest transcript density) to yellow (highest), set from 0.00 to 0.05. (F) Spatial transcript counts are shown as individual points, with low-quality transcripts (Q-score < 20) excluded to ensure accuracy. Scale bar = 200 μm in all panels.

**Fig. 6.**
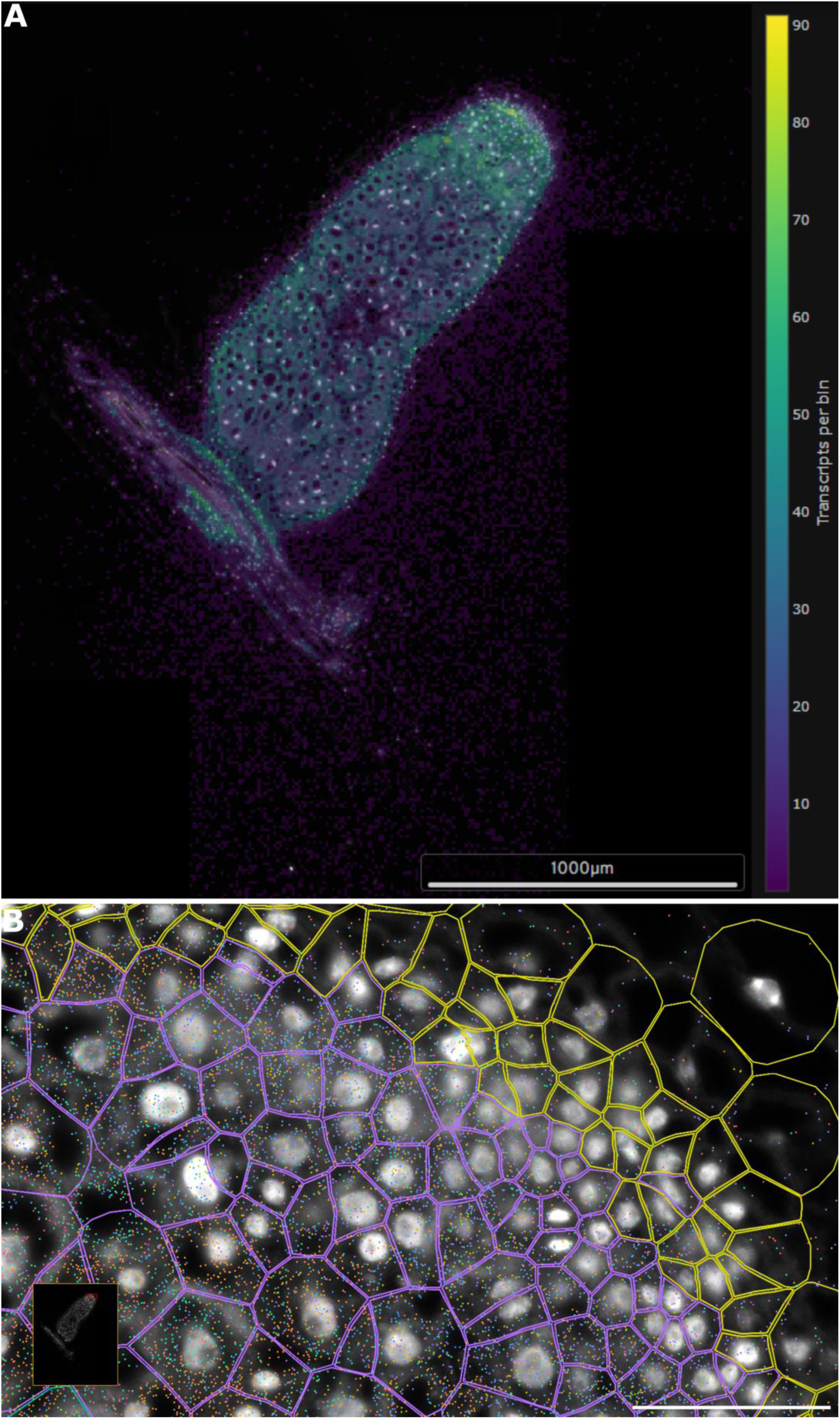
High-quality transcript detection and spatial resolution in mature *Medicago* nodules using a 50-gene custom panel. (A) DAPI-stained image of the analysed region overlaid with a heatmap representing the spatial distribution of high-quality decoded transcripts. The overlay is displayed with a bin size of 10 μm and can be adjusted for bin size and opacity. (B) Zoomed-in view of a subsection from panel A. All 50 transcripts are clearly detected, and unoccupied space remains, indicating potential to increase plexity with an add-on custom panel.

Note: If the lab already has a working H&E protocol, it can be used in place of this method; however, it should be tested on the tissue prior to use on the Xenium slide.

#### 3.5.2 Post-Xenium Quality Control Analysis

Once the Xenium raw data are received, the Xenium Analysis Summary file provides an overview of data quality. It is recommended to assess overall quality before interpreting the results. Ideally, if some of the target genes have well-characterized expression patterns, they can serve as internal validation markers. In our case, we used *MtNIN* gene, a well-studied gene in *Medicago*, known to have a defined expression pattern in mature nodules (*13*). This pattern has been previously validated using traditional in situ hybridisation (*13*). In our spatial transcriptomics dataset, NIN displayed the expected spatial expression with subcellular resolution (Fig. 5), supporting the success of the optimised protocol.

#### 3.5.3 Post-Xenium Confocal Imaging

The default Xenium cell segmentation method, designed for animal tissues, typically uses DAPI-stained nuclei with radial expansion—an approach that assumes spherical or elliptical cell geometries. However, plant cells exhibit a wide range of irregular, polygonal shapes and sizes, making this strategy suboptimal for plant tissues.

To improve segmentation, we recommend leveraging the intrinsic autofluorescence of plant cell walls, which can be captured using fluorescence lifetime imaging microscopy (FLIM) and tau gating on a confocal microscope. This provides a strong boundary signal, significantly enhancing segmentation accuracy. For plant species or tissue types with weak cell wall autofluorescence, cell wall-specific stains (e.g., Calcofluor White, Propidium Iodide, etc.) can be used to enhance boundary visibility in the images. This method allows more precise spatial mapping of transcripts to individual plant cells.

##### 3.5.3.1 Confocal Lambda-Lambda Scanning

Lambda scans acquire spectral emission data across a broad range of wavelengths, generating detailed fluorescence profiles. Certain confocal microscopes (e.g., Leica SP8) are equipped with white light lasers (WLL) for flexible excitation and prism-based detectors for tunable emission collection. These features are ideal for characterizing the specific autofluorescence profile of your plant tissue and identifying optimal excitation/emission settings for imaging.

*For detailed instructions, refer to your microscope manufacturer’s user manual for the lambda-lambda scan module*.

##### 3.5.3.2 Confocal FLIM Imaging

Once the optimal excitation and emission settings are identified, proceed with FLIM. This technique measures the fluorescence lifetime—the average time a fluorophore remains in an excited state before emitting a photon. FLIM generates images based on these lifetimes, with different colours representing different decay profiles.

Use fluorescence decay curve fitting to extract lifetime parameters for your tissue.

*Refer to your microscope’s user manual for FLIM-specific acquisition and analysis procedures*.

##### 3.5.3.3 Tau Gating and Tau Separation

While FLIM is highly informative, full-slide FLIM scanning can be time-consuming. Once fluorescence lifetime parameters are established, apply Tau Gating and Tau Separation using those parameters to efficiently filter unwanted autofluorescence and capture full-section images with enhanced cell wall boundary signals.

*Please consult your microscope manufacturer’s documentation for guidance on using Tau Gating/Tau Separation or equivalent functions*.

#### 3.5.4 Plant Cell Image Segmentation

Accurate segmentation of plant cells is essential for assigning transcripts to specific spatial compartments. While nuclei segmentation is readily handled by the default Xenium pipeline using the DAPI channel, plant tissues often require **cell wall-based segmentation** to better define true cell boundaries.

This section describes a workflow using **Cellpose**, a deep-learning-based image segmentation tool, to segment plant cells from fluorescence images, both from the DAPI channel typically provided in Xenium morphology.ome.tif files and cell wall autofluorescence images acquired by confocal microscope imaging.

##### 3.5.4.1 Image Segmentation Overview and Rationale

- **Nuclear segmentation** using the DAPI channel can be used when transcript assignment to nuclei is sufficient. The default Xenium segmentation is typically adequate.
- **Cell wall segmentation**, however, is preferred when precise cell boundary delineation is needed. This is especially relevant for spatial context analyses and is more challenging in plant tissues due to autofluorescence and irregular cell shapes.

Detailed code scripts and tutorials on installing and analysing are available on the GitHub repository: https://github.com/thiagomaf/PYcellsegmentation.

##### 3.5.4.2 Nuclei Segmentation (Default Approach)

Xenium’s DAPI channel is typically sufficient for nuclei segmentation, which is suitable when per-nuclear transcript assignment is adequate.

- **Input**: Xenium morphology.ome.tif file with DAPI channel.
- **Tool**: Cellpose version >= 4.0.5, using grayscale input image.
- Recommended settings:
- Estimate nuclear diameter in microns and convert to pixels using image metadata (µm/pixel).
- Use moderate resolution levels (e.g., Level 2) for faster processing.

This approach is robust and efficient, particularly for tissues where nuclei are evenly distributed and well-separated. However, it may not accurately reflect complex cell shapes in plant tissues.

##### 3.5.4.3 Cell Wall Segmentation (Recommended for Plants)

Segmentation based on cell wall staining provides more anatomically accurate boundaries, especially in tissues with large, irregularly shaped or densely packed cells.

- **Input**: Xenium morphology.ome.tif file with a fluorescent cell wall channel.
- **Tool**: Cellpose version >= 4.0.5, using grayscale input image.
- Workflow:

1. **Extract channel and resolution**: Use preprocessing scripts to extract the appropriate image pyramid level and isolate the cell wall stain channel.
2. **Tiling (if needed)**: Large images should be tiled with overlapping margins to reduce memory usage and improve segmentation accuracy.
3. Segmentation:

- Set an approximate cell diameter in pixels (based on tissue type).
- Tune Cellpose parameters such as flow_threshold, cellprob_threshold, and min_size for best results.
4. **Stitching**: If tiling was used, stitch segmented tiles into a full mask image.
- Advantages:

`o` Captures true plant cell geometry.
`o` Avoids assumptions based on nuclear position. Better suited for complex plant tissues.
- Quality control:

`o` Visualise results in Fiji, QuPath, or Napari by overlaying segmentation masks on the original image.
`o` Adjust parameters and rerun segmentation if boundaries are poorly defined or cells are merged.

##### 3.5.4.4 Segmentation Method Comparison and Selection

In plant tissues, selecting an optimal segmentation approach requires systematic evaluation, especially in the absence of manually annotated ground truth. We recommend a multi-metric strategy combining quantitative and qualitative assessments:

- Quantitative Metrics:

- Cell count and size distribution: Compare the number of detected cells and their size distributions across methods. Plant cells typically show characteristic size ranges for different tissue types (Table 16).
- Morphological features: Use shape descriptors such as aspect ratio and circularity to assess alignment with expected plant cell morphology (Table 16).
- Boundary quality: Evaluate the smoothness and continuity of cell boundaries, as plant cell walls should form continuous, well-defined borders.
- Qualitative Assessment:

- Biological plausibility: Assess whether the segmentation preserves known anatomical features and tissue organisation patterns (Fig. 7).
- Visual inspection: Overlay segmentation masks on original images to identify over-segmentation (artificially split cells) or under-segmentation (merged cells) (Fig. 7).
- Consistency: Compare results across similar tissue sections to evaluate method reproducibility.

**Fig. 7.**
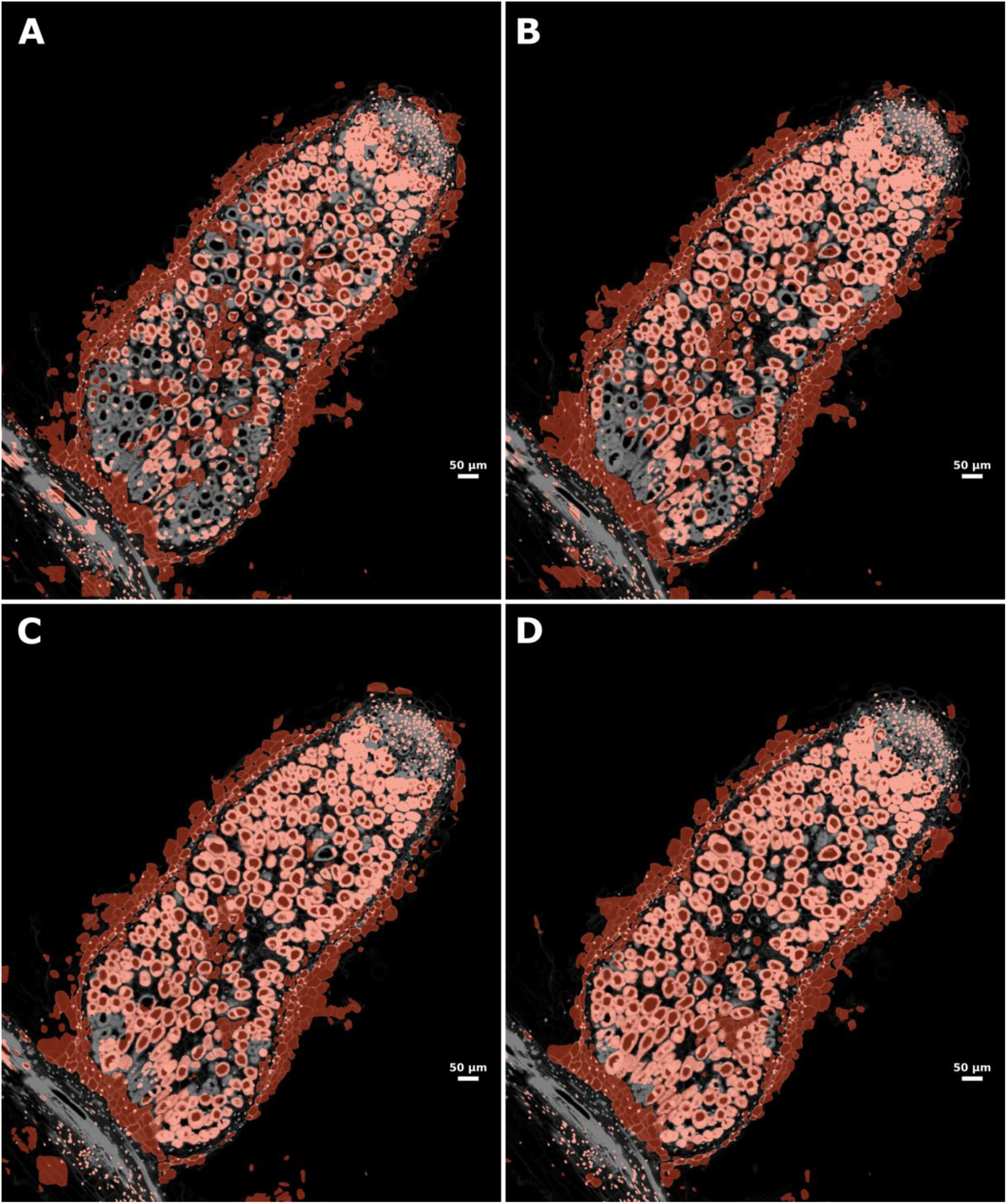
Cell segmentation using Cellpose. Panels show segmentation of sample B using different estimated cell diameters for Cellpose: (A) 6.38 µm, (B) 9.56 µm, (C) 12.75 µm, and (D) 15.94 µm. The background image shows the original DAPI-stained input. Red shading indicates segmented areas. The estimated cell diameter affects the results: 12.75 µm produces broader coverage overall, while 15.94 µm better captures larger infected cells within the fixation zone.

**Table 16.** Quantitative Comparison of Cell Segmentation Performance Across Different Cellpose Models. The table summarises key metrics derived from segmentation masks generated by various Cellpose parameter sets. ‘Diameter’ indicates the estimated cell diameter used as a model parameter. ‘Cell Count’ refers to the total number of unique cells identified, excluding the background. ‘Mean Area’ represents the average cell size. ‘Total Area’ indicates the cumulative area of all segmented cells.

| Sample | Diameter (μm) | Cell Count | Mean_Area (μm<sup>2</sup>) | Max_Area (μm<sup>2</sup>) | Total_Segmented Area (μm<sup>2</sup>) |
| --- | --- | --- | --- | --- | --- |
| **A** | 6.38 | 1912 | 338.09 | 2340.18 | 646425.81 |
|  | 9.56 | 1571 | 473.17 | 2606.78 | 743344.85 |
|  | 12.75 | 1449 | 531.77 | 3256.31 | 770539.75 |
|  | 15.94 | 1210 | 598.36 | 3954.24 | 724019.42 |
| **B** | 6.38 | 2112 | 312.31 | 2241.92 | 659608.54 |
|  | 9.56 | 1762 | 423.02 | 2457.22 | 745352.67 |
|  | 12.75 | 1604 | 494.18 | 3379.13 | 792667.03 |
|  | 15.94 | 1653 | 457.10 | 3912.34 | 755581.11 |
| **C** | 6.38 | 1818 | 305.23 | 1932.69 | 554902.40 |
|  | 9.56 | 1646 | 389.24 | 2748.39 | 640692.77 |
|  | 12.75 | 1436 | 471.70 | 2754.17 | 677363.26 |
|  | 15.94 | 1286 | 524.77 | 2939.85 | 674852.57 |

Comparative Analysis Results: To assess performance on *Medicago* nodule tissue, we applied three Cellpose models to serial sections and compared cell counts, aspect ratios, and circularities (Table 16). The systematic comparison approach can be adapted for other plant tissues by adjusting morphological expectations and evaluation criteria (Fig. 7).

#### 3.5.5 Post-segmentation Analysis and Transcript Mapping

After segmenting plant cells or nuclei, the next step is to assign individual transcripts to the segmented regions. This process enables the construction of a cell-by-gene expression matrix and downstream spatial analysis.

##### 3.5.5.1 Mapping Transcripts to Segmented Regions

- **Input requirements:**

`o` A segmentation mask image (e.g., from Cellpose), where each segmented cell or nucleus is labelled with a unique integer.
`o` A transcript coordinate file from Xenium output (e.g., .parquet or .csv), containing X/Y spatial positions and gene identity for each transcript.
- Transcript assignment:

`o` Each transcript is mapped to a segmented region by checking whether its spatial coordinates fall within a labeled area in the mask.
`o` Optionally, segmentation masks based on nuclei can be slightly expanded (e.g., by a few microns) to capture perinuclear transcripts if full cell boundaries are not available.
- Output:

`o` A mapped transcript table containing gene identity, cell/nucleus ID, and spatial coordinates.
`o` A cell-by-gene count matrix, which forms the foundation for downstream spatial transcriptomic analysis.

##### 3.5.5.2 Quality Control and Visualisation

- Segmentation QC: Overlay segmentation masks on the morphology image and check for accurate boundary definition.
- Mapping QC: Visualise mapped transcripts over segmentation results to verify correct assignment, especially at region edges or in densely packed tissues.
- Tools: Fiji, QuPath, and Napari can be used for both visual inspection and exploratory analysis.

## 4 Notes

1. Probe Design Considerations:

Gene selection is crucial for successful Xenium profiling. Avoid genes with very short transcripts (<500 bp), low expression, or extensive isoform overlap. Use reference expression datasets to prioritize genes showing tissue- or stage-specific enrichment.

2. Fixative Freshness:

Paraformaldehyde fixative should be prepared fresh each time for optimal crosslinking efficiency. Always cool the PFA solution on ice before adding glutaraldehyde to prevent degradation.

3. Vacuum Infiltration Efficiency:

Apply and release the vacuum gradually to prevent tissue damage. Monitor whether tissues sink—this indicates successful infiltration. Adjust vacuum time based on tissue density or species.

4. Tissue Orientation During Embedding:

Proper orientation is critical for downstream sectioning. When embedding root or nodule samples, place them vertically to maximise the number of longitudinal sections obtained per block.

5. Section Thickness Optimisation:

Xenium performs best on 5–10 µm sections. Thicker sections (>10 µm) may compromise probe penetration, while thinner sections may lead to tissue collapse, loss, or detachment during processing. If feasible, we highly recommend conducting a preliminary test run to evaluate how section thickness affects target transcript detection. Autofluorescence from the plant cell wall and the specific properties of the genes selected for probe design can both influence signal quality.

We suggest testing section thicknesses of 5 µm, 8 µm, and 10 µm. To guide your decision, evaluate metrics such as the high-quality transcript detection rate, the total number of high-quality transcripts detected, and visualized transcript density maps (Fig. 6). For *Medicago* nodules, we found that 8 µm and 10 µm sections yielded significantly better results than 5 µm in terms of target detection efficiency.

6. Floating Time Optimisation:

Optimal floating time can vary for different tissue types and should be tested prior to Xenium sectioning. Shorter times can lead to wrinkles in the section, whilst longer times can allow the tissue to stretch too far. 5 – 10 seconds was found to be optimal for M*edicago* tissue, as longer times resulted in over-stretching and splitting of the vasculature tissue.

1. Ribonuclease-Free Handling:

Always use autoclaved or filtered MQ water, RNAse-free tips, and clean slides to reduce contamination. Use gloves and avoid talking directly over open slides when mounting tissue.

2. Slide Storage Before Hybridisation:

Store dried slides in a dust-free, low-humidity environment such as a desiccator. Slides can be stored up to 4 weeks, but longer storage may reduce signal strength.

3. Autofluorescence Quenching: Plant tissues, especially root nodules, often exhibit high autofluorescence. Always include quenching steps prior to imaging to improve the signal-to-noise ratio.

4. Quality Control (QC) with H&E or DAPI: Before proceeding with expensive hybridisation steps, perform H&E or DAPI staining to ensure tissue integrity and RNA preservation (Fig. 5). This step also helps in adjusting the section thickness.

5. Tissue Compression During Sectioning: If sections appear wrinkled or compressed, ensure the microtome blade is sharp and the paraffin block is at the correct cutting temperature. Chilling the block slightly before sectioning can help.

## 5 Results

To evaluate the performance of our optimised Xenium in situ workflow on plant tissues, we applied the protocol to formalin-fixed paraffin-embedded (FFPE) sections of mature *Medicago truncatula* root nodules harvested 28 days post-inoculation. Using an 8 μm section and a 50-gene custom panel, we successfully detected spatially resolved transcript expression across major nodule regions (Fig. 5). Clear nuclear staining and tissue autofluorescence (Fig. 5A) confirmed good preservation, and spatial expression patterns of marker genes (Fig. 5B) aligned with known nodule zonation. Post-Xenium H&E staining (Fig. 5C) verified the maintenance of tissue morphology. *NODULE INCEPTION* (*MtNIN*) expression with high-confidence transcripts (Q-score ≥ 20) was spatially enriched in the infected zone (Fig. 5D–F).

The 50-gene panel demonstrated strong spatial resolution and transcript detection capacity (Fig. 6). High-quality decoded transcripts were evenly distributed across the tissue, with individual transcripts clearly localised and ample physical space remaining for additional probe sets. These results suggest that our base panel provides robust detection with capacity for multiplexing via add-on panels.

Finally, we tested cell segmentation using Cellpose with varying diameter estimates (Fig. 7). A diameter of 15.94 μm yielded optimal segmentation of larger infected cells within the fixation zone, while smaller diameters provided more general coverage across nodule cell types. This flexible segmentation approach enables customisable cell boundary identification to match the tissue context.

Together, these results validate the effectiveness and reproducibility of our Xenium in situ protocol for high-resolution spatial transcriptomics in *Medicago* nodules, enabling detailed exploration of cell-type-specific gene expression in plant developmental contexts.

## Author Contributions

MYJ initiated the project and wrote the abstract and introduction. MYJ, JH, AD, TAM, CX, and AMP collaboratively drafted the initial version of the methods section. All authors reviewed and edited the manuscript and approved the final version for publication.

MYJ curated and selected the gene list and conducted probe design with assistance from the 10x Genomics bioinformatics team. CX and MYJ identified orthologs between *Medicago truncatula* A17 and R108 accessions. This ortholog information was used to guide conserved probe design, carried out by MYJ in collaboration with the 10x Genomics team.

MYJ designed the experiment and conducted sample harvesting and FFPE block preparation. JH performed tissue sectioning in consultation with MYJ to ensure capture of target regions. JH and the histology team conducted deparaffinization to prepare slides for the Xenium workflow. AD and AMP performed the 10x Xenium workflow and generated raw Xenium data. JH and the histology team conducted post-Xenium H&E staining. MYJ performed post-Xenium confocal scanning. TAM conducted cell segmentation analysis. MYJ carried out gene-of-interest analysis and quality control assessment.

## Acknowledgements

M.Y.J. is supported by grants made to the University of Cambridge by the Bill and Melinda Gates Foundation and the UK Foreign, Commonwealth and Development Office (INV-006871) and Bill & Melinda Gates Agricultural Innovations (INV-57461) known as the Enabling

Nutrient Symbioses in Agriculture (ENSA) project. We acknowledge the support from the 10x Genomics team for their assistance with custom Xenium panel design and data interpretation. We also thank the Histopathology/ISH Core Facility, Cancer Research UK—Cambridge Institute, for their technical support with H&E staining and bright-field scanning of slides. Additional thanks to the Cancer Research UK Cambridge Institute Genomics Cores for their support with various aspects of this study, and to Rachel Barnes for performing Xenium workflows and generating Xenium raw data for the 50-gene runs. Finally, we thank Eli Marable for his valuable feedback, which helped enhance readability and clarity.

